# Functional characterization of 5’ UTR *cis*-acting sequence elements that modulate translational efficiency in *P. falciparum* and humans

**DOI:** 10.1101/2021.09.07.459299

**Authors:** Valentina E. Garcia, Rebekah Dial, Joseph L. DeRisi

## Abstract

**Background:** The eukaryotic parasite *Plasmodium falciparum* causes millions of malarial infections annually while drug resistance to common antimalarials is further confounding eradication efforts. Translation is an attractive therapeutic target that will benefit from a deeper mechanistic understanding. As the rate limiting step of translation, initiation is a primary driver of translational efficiency. It is a complex process regulated by both *cis* and *trans* acting factors, providing numerous potential targets. Relative to model organisms and humans, *P. falciparum* mRNAs feature unusual 5’ untranslated regions suggesting *cis*-acting sequence complexity in this parasite may act to tune levels of protein synthesis through their effects on translational efficiency.

**Methods:** Here, we deployed *in vitro* translation to compare the role of *cis*-acting regulatory sequences in *P. falciparum* and humans. Using parasite mRNAs with high or low translational efficiency, the presence, position, and termination status of upstream “AUG”s, in addition to the base composition of the 5’ untranslated regions, were characterized.

**Results:** The density of upstream “AUG”s differed significantly among the most and least efficiently translated genes in *P. falciparum*, as did the average “GC” content of the 5’ untranslated regions. Using exemplars from highly translated and poorly translated mRNAs, multiple putative upstream elements were interrogated for impact on translational efficiency. Upstream “AUG”s were found to repress translation to varying degrees, depending on their position and context, while combinations of upstream “AUG”s had nonadditive effects. The base composition of the 5’ untranslated regions also impacted translation, but to a lesser degree. Surprisingly, the effects of *cis*-acting sequences were remarkably conserved between *P. falciparum* and humans.

**Conclusion:** While translational regulation is inherently complex, this work contributes toward a more comprehensive understanding of parasite and human translational regulation by examining the impact of discrete *cis*-acting features, acting alone or in context.

## Background

As the primary cause of severe malaria, *Plasmodium falciparum* remains a major global health threat. In 2018, approximately 228 million cases of malaria led to 405,000 deaths, primarily of children under the age of 5 [1]. Control and eradication of *P. falciparum* is complicated by widespread or emerging drug resistance to all common antimalarial drugs [2–4]. To circumvent drug resistance, targeted therapeutic development has the potential to generate novel antimalarials with unique mechanisms of action. Unfortunately, targeted development is hindered by an incomplete understanding of the basic molecular processes of *P. falciparum* and how they differ from human biology.

Recently, translation has emerged as a potentially druggable pathway [5–7]. While no clinically approved antimalarials target cytoplasmic translation [5], there are promising new candidates to distinct translational mechanisms. For example, there is a growing number of compounds targeting tRNA synthetases [8, 9], M5717 (formerly DDD107498) is currently in human trials and inhibits eukaryotic elongation factor 2 [6, 8], and MMV008270 has been shown to selectively inhibit parasite translation through an unknown mechanism of action [10]. Currently no candidates are known to target translation initiation.

Eukaryotic translation initiation determines the rate of translation of a given mRNA, referred to as the translational efficiency (TE) [11, 12]. Initiation at the proper translation start site (typically an “AUG” start codon) relies on interactions between the start codon and the local sequence context (the Kozak sequence) with the initiator Met-tRNA and other initiation factors [13–15]. TE can additionally be regulated by *cis*-acting sequence elements throughout the 5’ untranslated region (5’ UTR), the sequence proceeding the translation start site. In particular, upstream “AUG”s (uAUGs) are commonly observed regulatory features that are divided into two groups: those that initiate open reading frames that extend beyond the translation initiation site, and those that are terminated, meaning they form upstream open reading frames (uORFs) by having an in-frame stop site proceeding the protein coding region [16–18]. These *cis*-acting regulatory elements lower TE through many potential mechanisms including by initiating translation out of frame from the downstream ORF, by adding long amino acid extensions at the N-terminus, or by sequestering ribosomes within the 5’ UTRs [19–21].

A well-documented example of uAUG/uORF driven regulation is GCN4 in *Saccharomyces cerevisiae*. The 5’ UTR of GCN4 contains four short uORFs that themselves are differentially translated under conditions of stress. Based on the availability of translation initiation factors, the uORFs modulate the translation rate of the primary protein coding region to fit the organisms current nutrient conditions [22, 23]. While this example is deeply understood, it is not broadly generalizable, and the rules by which such sequences exert influence on TE remain challenging to describe even for the most studied of eukaryotes. For example, numerous variables have been identified in other contexts that modulate the effect of uAUGs and uORFs, including the Kozak sequence of the uAUG itself and the reading frame relative to the translational start site [19, 24].

Studies of *P. falciparum* have confirmed that it possesses the expected eukaryotic cap-binding factors required for cap-dependent translation initiation [25, 26]. Additionally, gene specific studies show that uAUGs and uORFs can repress translation in *P. falciparum* and that the Kozak sequence of uAUGs along with uORF length may modulate their effect on TE [27, 28]. This is particularly intriguing since *P. falciparum* has repeatedly been shown to have unusually long 5’ UTRs containing many uAUGs [18,29,30]. Together this suggests that multiple *cis*-acting factors within the 5’ UTRs of *P. falciparum* could act broadly to tune TE throughout the normal lifecycle, as opposed to regulating specific genes under extreme conditions, such as with GCN4 regulation. However, extensive ribosome profiling from our lab revealed that transcription and translation rates are highly correlated throughout the intraerythrocytic life cycle with less than 10% of the transcriptome being under significant translational control [18]. Ribosome profiling also showed that the presence of uAUGs and uORFs did not appear to correlate with TE, which is in contrast to model organisms and classic paradigms like yeast GCN4. Together this highlights that it remains difficult to predict how *cis*-acting sequences within a given 5’ UTR will affect TE, especially in disparate eukaryotic species.

Here, we sought to understand the interconnected effects of 5’ UTR *cis*-acting regulatory elements with respect to TE in both *P. falciparum* and human cells through a highly reductionist approach. To do so, we deployed an *in vitro* translation assay for *P. falciparum* and developed an equivalent assay for human K562 cells. Using a pair of naturally occurring *P. falciparum* 5’ UTRs with differing TEs, the individual contributions of the sequence context, positionality, and termination status of uAUGs, along with the base composition of the 5’ UTR to TE, were systematically dissected to understand their contributions, in isolation and in combination. Together these data present a complex portrait of interacting elements within 5’ UTRs that directly influence TE, most of which are similar in both *P. falciparum* and human.

## Methods

### Identifying characteristics associated with high and low TE from the 5’ UTRs of *P. falciparum*

The ribosome profiling and mRNA sequencing data from the late trophozoite stage generated by Caro and Ahyong *et. al.* 2014 [18] were filtered for an abundance above 32 reads per million, a TE greater than zero, and a predicted 5’ UTR length above 175 nucleotides. Additionally, 30 genes that are not included in the PlasmoDB-28 *P. falciparum* 3D7 gene annotations were removed. This resulted in a data set containing 2088 genes (Additional File 1). The 5’ UTR sequences were determined using the PlasmoDB-28 *P. falciparum* 3D7 genome. Sequence analysis was done using Python, K.S. tests were done using the Python SciPy package, and the data for Figure 1 was graphed using the Python Matplotlib package.

**Figure 1:**
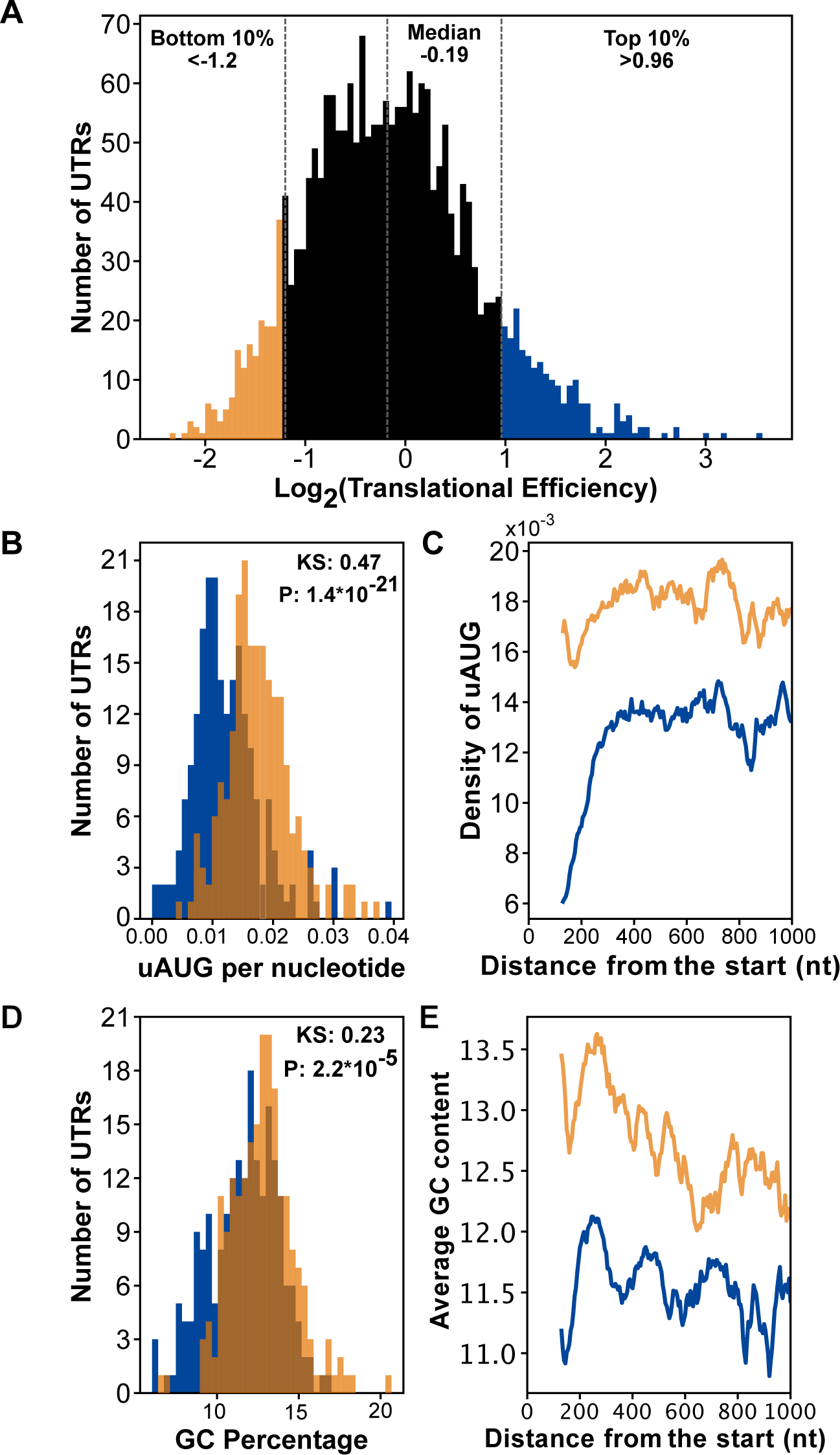
Comparison of features within the 5’ UTRs of genes in the bottom 10% (n=209) and top 10% (n=209) of TEs in the late trophozoite stage using data from Caro and Ahyong *et. al.* 2014 [18]. a) A histogram of the log_2_(TE)s of genes expressed in the late trophozoite stage included in subsequent analysis. The vertical dotted lines indicate the bottom 10% (yellow) and top 10% (blue) of 5’ UTRs. b) The number of uAUGs normalized to the length of the 5’ UTRs in the bottom 10% (yellow) and top 10% (blue) of TEs in the late trophozoite stage. The two distributions are statistically distinct, KS test-statistic 0.47, p-value 1.4*10^-21^. c) The average number of uAUGs in the 5’ UTRs within a 130-nucleotide window sliding by 5 nucleotides up to 1000 nucleotides of the bottom 10% (yellow) and top 10% blue. d) The distribution of GC content in the bottom 10% (yellow) and top 10% (blue) of TEs in the late trophozoite stage are statistically distinct, KS test statistic comparison of the two: 0.23 p-value 2.24*10^-5^. e) The average GC content within a 130-nucleotides sliding window moving 5 nucleotides up to 1000 nucleotides from the translation start site. Bottom 10% (yellow) and top 10% blue.

### Cloning length variations of PF3D7_1411400 and PF3D7_1428300 5’ UTRs

The first step to generating the constructs used here was to create a Puc118-NanoLuc construct without a 5’ UTR (Additional File 2). Using In-fusion cloning the 5’ UTR and firefly luciferase enzyme from the EBA175-Firefly plasmid used previously [5, 10] were replaced with the NanoLuc Luciferase (Promega) coding sequence. The plasmid generated, called P16, consists of: Puc118 backbone with a T7 promotor proceeding the NanoLuc Luciferase protein coding sequence followed by the 3’ UTR from PF_HRP2.

To create the varying length 5’ UTR constructs, the 5’ UTR sequences of PF3D7_1411400 and PF3D7_1428300 were amplified from *P. falciparum* W2 strain gDNA using Kapa 2G Robust DNA polymerase (Roche KK5024) with primers containing overhangs with the T7 promoter (forward primer) or NanoLuc (reverse primer). The P16 plasmid was amplified using Phusion polymerase (NEB M0530S) for the backbone (forward primer: ATGGTCTTCACACTCGAAGATTTC, reverse primer: CCTATAGTGAGTCGTATTAGAATTCG). The inserts and backbone were purified using a Zymo DNA Clean and Concentrator-5 (Zymo Research D4013). In-fusion reactions were performed per the In-fusion Cloning Kit (Takara 638918) instructions and reactions were transformed into Stellar Competent Cells (Takara 636766).

### Cloning 5’ UTR 130 nucleotide constructs

To generate the 130 nucleotide 5’ UTR constructs, long oligos containing an EcoRI-HF cut site, the T7 promoter, the desired 5’ UTR sequence, and a priming sequence to NanoLuc were purchased from Integrated DNA Technologies (forward primer: TGATTACGAATTCTAATACGACTCACTATAGG-desired 5’ UTR - ATGGTCTTCACACTCGAAGATTTC).

The P16 plasmid was used as a template for PCR using Kapa 2G Robust with the reverse primer binding just after the BamHI-HF restriction site in Puc118 (reverse primer: CTGCAGGTCGACTCTAGA). PCR products were run on a 1% agarose gel to check the product size and purified using Zymo DNA Clean and Concentrator-5 (Zymo Research D4013). To create the cloning insert, purified PCR product was digested with EcoRI-HF and BamHI-HF at 37°C for 1.5 hours and purified again using a Zymo DNA Clean and Concentrator-5 (Zymo Research D4013). For the cloning backbone, P16 was digested with EcoRI-HF and BamHI-HF at room temperature overnight (∼12 hours), run on a 1% agarose gel, and gel extracted with the Zymoclean Gel DNA Recovery Kit (Zymo Research D4008). The insert and backbone were ligated using T4 DNA ligase (NEB M0202S) at room temperature for 30 mins and heat inactivated at 65°C for 10 mins. After heat inactivation, the reaction was transformed into Stellar Competent Cells (Takara 636766). All constructs were sequence verified. The sequences for all the 5’ UTRs evaluated can be found in Additional File 3.

### Generating reporter RNA for *in vitro* translation

All mRNA generating plasmids were digested with PvuII-HF (NEB R[Δ4]151L) and ApaLI-HF (NEB R0507L) at 37°C for 3 hours. After digestion, templates were run on a 1% agarose gel to confirm cutting and the reactions were purified with Zymo DNA Clean and Concentrator-5 (Zymo Research D4013). 1ug of linearized template was used in a 100uL T7 RNA Polymerase (purified in house) reaction that was incubated at 37°C for three hours. After T7 reactions were complete, 15uL TurboDNAse (ThermoFisher Scientific AM2238) was added, and reactions were incubated at 37°C for 15 minutes. The RNA was then purified using a Zymo RNA Clean and Concentrator-25 Kit (Zymo Research R1017). Eluted RNA was measured using the Qubit RNA HS Assay Kit (Thermo Fisher Scientific Q32852) then capped following the protocol for the Vaccinia Capping System (NEB M2080S) and purified one last time using Zymo RNA Clean and Concentrator-5 (Zymo Research R1013). Capped RNA concentrations were measured using the Qubit RNA HS Assay Kit (Thermo Fisher Scientific Q32852). Final RNA was diluted to 0.25pmoles/ul for use in the *in vitro* translation assays.

For the comparing capped veruses uncapped mRNA, uncapped RNA was incubated for 5 minutes at 65°C to match the treatment of capped RNAs. The same RNA that was used in the vaccinia capping reaction was directly compared to the post-cap RNA.

### Generating *P. falciparum in vitro* translation lysates

*P. falciparum* W2 strain (MRA-157) from MR4 was grown in human erythrocytes at 2% hematocrit in RPMIc medium (RPMI 1640 media supplemented with 0.25% Albumax II (GIBCOLife Technologies), 2 g/L sodium bicarbonate, 0.1 mM hypoxanthine, 25 mM HEPES (Ph 7.4), and 50μg/L gentamicin), at 37°C, 5%O2, and 5%CO2. Cultures were maintained at 2-5% parasitemia.

In depth step-by-step protocols for lysate generation have been previously published [5]. In summary, cultures were synchronized twice using 5% sorbitol six hours apart. Once cultures recovered to 10% parasitemia, they were used to seed two 500mL hyperflasks (Corning 10031). When cultures reached the late trophozoite stage at 10−20% parasitemia, the cultures were centrifuged for 5 min at 1500g at room temperature with no break, the supernatant was removed, and 0.025–0.05% final saponin (exact amount determined by optimization of each batch of saponin) in Buffer A (20 mM HEPES pH8.0, 2mM Mg(OAc)_2_, 120mM KOAc) was added. Saponin lysed cultures were centrifuged at 4°C at 10,000g for 10 min in a Beckman Coulter J26XPI. Pellets were washed twice with buffer A with centrifuging between each wash and then were re-suspended in an equal volume to the pellet of BufferB2 (20 mM HEPES pH8.0,100 mM KOAc, 0.75mMMg(OAC)_2_, 2mMDTT,20% glycerol,1XEDTA-free protease inhibitor cocktail (Roche)), flash frozen, and stored in -80°C. Frozen pellets were then thawed at 4°C and lysed by passing them through a cell homogenizer containing a 4μm-clearance ball bearing (Isobiotec, Germany) 20 times by hand or using a custom build machine [31]. The whole-cell lysate was then centrifuged at 4°C at 16,000g for 10 min and the supernatant was flash frozen and stored at -80°C. The experiments performed here used a pool of lysates from multiple different harvests that were each individually tested for a minimal activity of 10^4^ using a high expression RNA containing NanoLuc (A[WT]) and the Promega Nano-Glo Luciferase assay system (Promega N1110). Pooled lysates were then optimized for the needed amount of Mg(OAc)_2_ and an optimal incubation time at 37°C, in this case 3mM final concentration Mg(OAc)_2_ and 57 minutes.

### Generating K562 *in vitro* translation lysates

K562 suspension cells were cultured in RPMI 1640 media supplemented with 10% fetal bovine serum, 10mM Hepes (pH 7.2-7.5), and 0.5mg/mL Penicillin-Streptomycin-Glutamine. Cultures were maintained by splitting to 10^5^ cells/mL and were counted using a BD Accuri.

When cells reached 10^6^ cells/mL, the cultures were centrifuged for 5 min at 1500g at room temperature and the supernatant was removed. Pellets were washed twice with buffer A with centrifuging at 1500g at 4°C between each wash. Finally, pellets were re-suspended in an equal volume of Buffer B2 and flash frozen in liquid nitrogen. Cell lysates were generated from the frozen pellets using the same methodology as *P. falciparum* lysates, but with the cell homogenizer containing a 12 μm-clearance. Lysates that produced over 10^4^ luminescence units sing a high expression RNA containing NanoLuc Luciferase (A[WT]) and the Promega Nano-Glo Luciferase assay system (Promega N1110) in preliminary tests were pooled and optimized for the needed amount of Mg(OAc)_2_ and incubation time using A[WT] mRNA, in this case 1.5mM final concentration Mg(OAc)_2_ and 12 minutes.

### *In vitro* translation protocol

*In vitro* translation reactions for *P. falciparum* and K562 lysates were set up identically. 3uL of buffer B2 and 2uL of 0.25pmole/uL RNA were placed into 384-well plates. A master mix of 3.5uL lysate with 0.5uL 100uM complete amino acid mix (Promega L4461) and 1uL 10x translation buffer (20mM Hepes pH 8, 75mM KoAc, 2mM DTT, 5mM ATP, 1mM GTP 200mM creatine phosphate, 2ug/ul Creatine kinase, and the pre-determined concentration for each lysate pool of Mg(OAc)) was added to each well. Reactions are then incubated for the pre-determined amount of time at 37°C, then placed on ice to stop the reactions. 8uL of reaction was mixed with 8uL of Nano-Glo buffer/substrate mix following the Nano-Glo Luciferase Assay System (Promega N1110) instructions. Luminescence was measured on a Promega GloMax Plate Reader (Promega TM297) with a 6 second integration time.

### Analysis

#### Experimental TEs

All experiments were performed three separate times in triplicate, for a total of 9 values per mRNA tested (except for the capped and uncapped experiment which was done 3-4 times in duplicate). For each separate experiment, new mRNA was generated and capped. For the figures, each value was normalized to the mean of the triplicates from each separate run. All raw values and normalized values can be found in Additional File 4. The fold differences were log_2_ transformed and then used to calculate the mean and SEM. Graphs for the figures were made using a custom Python/Postscript script (Additional File 5).

#### Predicted activity of multiple uAUGs

The percent repression of each uAUG individually was calculated by determining the percent of R[Δ1:Δ2:Δ3:Δ4] (the “maximum signal”). For the predictions, the percent repression of each uAUG in the model were multiplied together.

#### Predicted Secondary Structures

To evaluate for secondary structure, the ΔG of 30 nucleotide stretches of the 5’ UTR tiled with a 5 nucleotide separation was generated using RNAfold [32]. The predicted ΔG were then plotted using GraphPad Prism Software.

## Results

### Identifying putative *cis*-acting elements within the 5’ UTRs of *P. falciparum* that differ between genes with high and low TE

To identify putative *cis*-acting sequences that regulate TE in *P. falciparum*, the ribosome profiling and mRNA sequencing data generated by Caro and Ahyong *et. al.* [18] was re-analyzed by comparing the 5’ UTR sequences of genes in the bottom 10% and top 10% of TEs during the late trophozoite stage (Figure 1A). Features within the 5’ UTRs were quantified, and the distributions from each set were compared. While the distributions of 5’ UTR length were not statistically distinct (K.S. test p=0.10 Supplemental Figure 1A), the distributions of uAUG frequency differed significantly and appeared distinctly separated when normalized to 5’ UTR length with lower TE genes tending to contain more uAUGs (K.S. test p= 3.36*10^9^ and -21 p=1.1*10^-21^ respectively) (Supplemental Figure 1B, Figure 1B). This trend appeared to be most distinct closest to the protein coding region (Figure 1C).

Additionally, the distributions of GC content statistically differed between 5’ UTRs with low and high TE (K.S. test p=2.24*10^-3^) (Figure 1D). The positional effect followed a similar trend with repressed genes on average having a higher GC content, especially near the translational start site (Figure 1E). Together, this retrospective bioinformatic analysis suggested that these two features should be further investigated for their role in influencing TE with particular attention placed on the sequence region proximal to the translation start site.

### Evaluating *P. falciparum* and human K562 *in vitro* translation assays for measuring the effect of 5’ UTRs on TE

To investigate the role of *cis*-acting elements within 5’ UTRs, an *in vitro* translation assay previously developed for identifying translation inhibitors against *P. falciparum* [5, 10] was adapted using both *P. falciparum* W2 and *H. sapiens* K562 cellular extracts. To validate and optimize the platform for this purpose, two mRNAs transcribed in late trophozoites with significantly different TEs were identified, PF3D7_1411400 (a plastid replication-repair enzyme) representing a translationally repressed mRNA from the bottom 10% of TEs and PF3D7_1428300 (a proliferation-associated protein) representing a high translation mRNA from the top 10% of TEs. These two genes were chosen for their relatively similar 5’ UTR lengths and other properties (Figure 2A and B). The full length 5’ UTRs of both genes (Figure 2A) were cloned into a reporter construct driving expression of a luciferase enzyme and were evaluated for their effect on TE.

**Figure 2:**
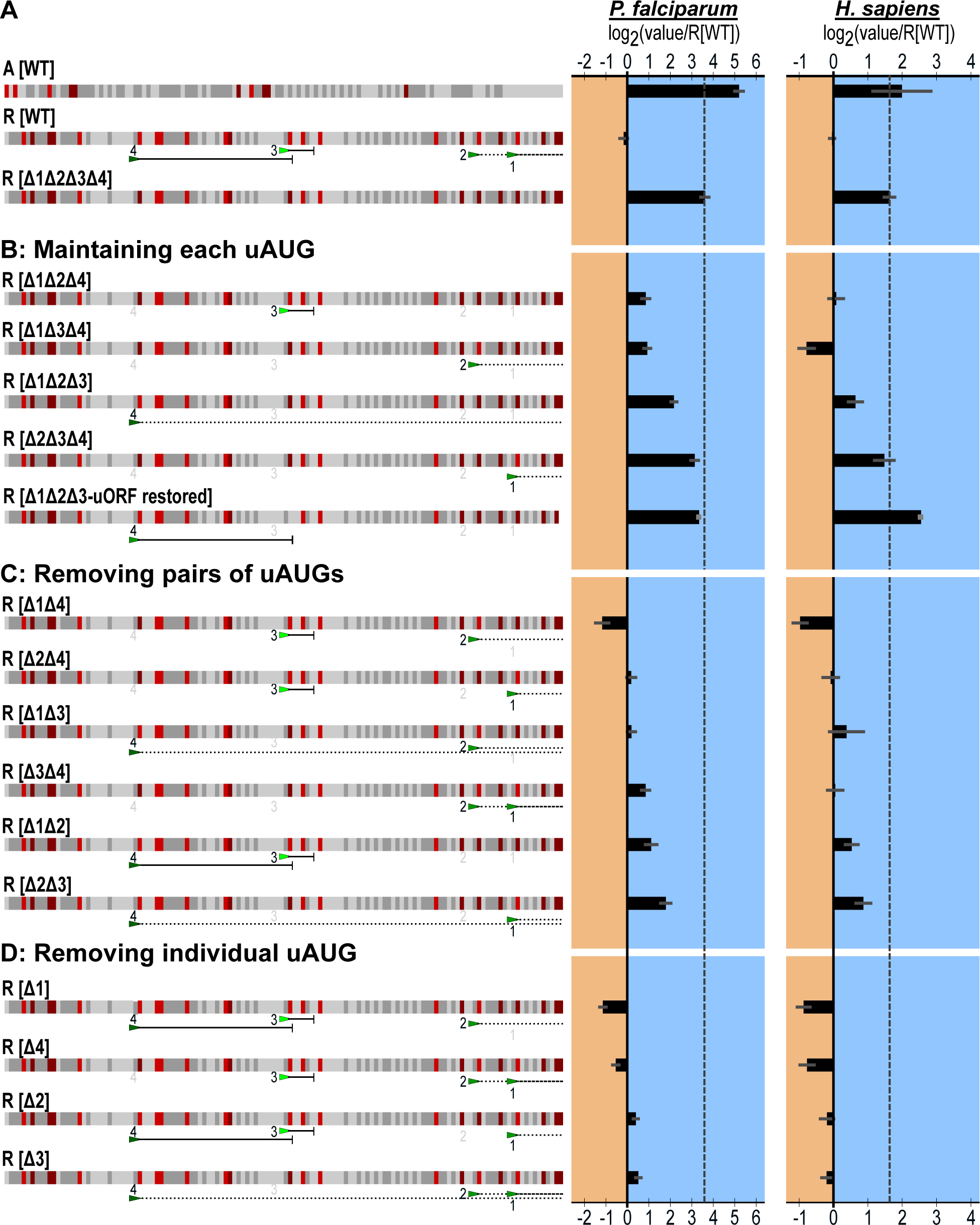
130 nucleotides of the 5’ UTR from a translationally active (PF3D7_1428300) and repressed gene (PF3D7_1411400) were sufficient to drive distinct TE. a) The diagramed sequence of the full length 5’ UTRs from active PF3D7_1428300 and repressed PF3D7_1411400. uAUGs are marked by green triangles with the different shades representing the three frames. The numbers between the two construct diagrams mark distance from the protein coding region. b) The lengths, uAUG count, uORF count, GC content, and translational efficiency (TE) for the chosen 5’ UTRs and genes obtained from the previously published ribosome profiling and mRNA sequencing [18] with the raw luminescence signal (PF RLU) produced by these 5’ UTRs driving NanoLuc (Promega) expression using *P. falciparum in vitro* translation. c) Log_10_(luminescence) from NanoLuc produced by 5’ UTRs of decreasing length in *P. falciparum* lysates (red) and K562 lysates (grey). The different lengths were generated by shorting the 5’ UTRs from the 5’ end. d) Sequence comparison of the 130 nucleotides closest to the protein coding region of the 5’ UTRs from PF3D7_1411400 (R[WT]) and PF3D7_1428300 (A[WT]). The four uAUGs in R[WT] are labeled with the green triangles. uAUGs without in-frame stops are followed by a dotted line while uORF forming uAUGs are followed by a solid line with the stop is marked by a vertical line. The four uAUGs are labeled 1-4 based on their distance from the protein coding start site.

The 5’ UTR of PF3D7_1411400 is 730 nucleotides long, contains 15 uAUGs (13 form uORFs), and is 11.0% GC (Figure 2B). Using the data of Caro *et. al.* [18], the RNA abundance was measured to be 63.66 reads per million and the log_2_(TE) was -1.94. The 5’ UTR of PF3D7_1428300 is 775 nucleotides long, contains 10 uAUGs (all of which form uORFs), and is 9.3% GC (Figure 2B). The abundance for the RNA was measured to be 522.93 reads per million and the log_2_(TE) was 1.75. Thus, the TE of the active gene is 12.2-fold higher than that of the repressed gene by ribosome profiling. In the *P. falciparum in vitro* translation assay, which effectively removes any influence from differential expression levels, the signal produced by the activating 5’ UTR was 24.5-fold higher than the signal from the repressive 5’ UTR (Figure 2B). In the K562 *in vitro* translation assay, the 5’ UTR from the active gene also out-performed that of the repressed gene by 5.3-fold (Figure 2C). Both *in vitro* translation assays recapitulated the difference in TE that was observed *in vivo*, albeit with different absolute magnitudes.

As noted above, the 5’ UTR analysis of the ribosome profiling data suggested that differences between high and low TE 5’ UTRs appeared to be exaggerated closer to the translation start site. To investigate this while reducing the search space for *cis*-acting elements, each of the 5’ UTRs was progressively trimmed from the 5’ end (Figure 2C). In *P. falciparum* lysates, shortening the activating 5’ UTR to 549 nucleotides increased translation 4.2-fold, and reducing the UTR to 130 nucleotides further increased translation 1.9-fold, for a 7.9-fold total increase. Reducing the repressive 5’ UTR to 339 nucleotides similarly increased translation 3.15-fold, but further reduction to 130 nucleotides resulted in no additional increases in *P. falciparum*. Similarly, in human K562 lysates, trimming of the 5’ UTRs resulted in an overall increase in translation for both 5’ UTRs and increased the TE differential between the two (Figure 2B).

While trimming both 5’ UTRs increased their respective translation, the differential between the activating and repressive UTRs was magnified. At 130 nucleotides, the activating 5’ UTR outperformed the repressive 5’ UTR by 64-fold (Figure 2B), which had the added benefit of increasing the dynamic range between constructs. Hence forth, the minimal 130 nucleotide sequences were used as the platform for further dissection of *cis*-acting sequences and all subsequent 5’ UTRs evaluated were 130 nucleotides. The activating 130 nucleotide 5’ UTR derived from PF3D7_1428300 is denoted as A[WT] and the repressive 130 nucleotide 5’ UTR from PF3D7_1411400 is denoted as R[WT]. Reflective of the distinct distributions in uAUG abundance and GC abundance, R[WT] is 16.9% GC and contains four uAUGs, numbered 1-4 based on distance from the translation start site. uAUGs 1 and 2 do not form uORFs and are in the +1-frame relative to the reporter gene starting at -13 and -22 nucleotides, while uAUGs 3 and 4 both form uORFs at -66 and -101 nucleotides. A[WT] is 7.7% GC and contains no upstream “AUG”s (Figure 2D).

All the RNAs used herein were capped using Vaccinia Capping Enzyme (NEB M2080S). To verify that both lysates were sensitive to capping, capped and uncapped versions of the full length 5’ UTRs and the 130 nucleotide 5’ UTRs were compared (Supplemental Figure 2). Both lysates were sensitive to capping, with capped RNAs generally generating more luminescence (up to a 21.7-fold increase in *P. falciparum* and 7.1 in K562 with full length 1429300), especially in *P. falciparum* lysates. Additionally, in K562 lysates, uncapped RNAs with the full length 5’ UTRs generated a more variable signal than capped RNAs. To promote scanning initiation, increase luminescence signal, and reduce noise, all further experiments in this study utilized capped RNA.

### Measurement of both independent and combined effects of uAUGs on translational repression

The combined effect of the four uAUGs in R[WT] was first evaluated by mutating all four to “AUC”, denoted R[Δ1Δ2Δ3Δ4]. Conversion of all four alleviated repression by over 1000% in *P. falciparum*, and 337% in human lysates (Figure 3A). If each uAUG equally contributed toward repression, the expected result of maintaining any single uAUG would be a consistent relief from repression relative to R[WT]. However, individually maintaining each of the four uAUGs yielded significantly different degrees of translation (Figure 3B), ranging from a modest 2-fold increase with uAUG-3 alone (R[Δ1Δ2Δ4]) to a nearly 10-fold increase with uAUG-1 alone (R[Δ2Δ3Δ4]), indicating unequal contributions towards the overall level of repression. For K562 extracts, the results were similar, although uAUG-2 alone (R[Δ1Δ3Δ4]) was the most repressive of the set, being even more so than the wild-type construct. Since uAUG-4 forms a uORF whose stop site overlaps with uAUG-3 and was eliminated by making uAUG-3 into “AUC”, uAUG-4 with a restored uORF was also evaluated (R [Δ1Δ2Δ3-uORF restored]). With the uORF restored, uAUG-4 confers minimal or no translational repression. These data demonstrate that each of the individual uAUGs in isolation possess differing repressive activities with respect to translation.

**Figure 3:**
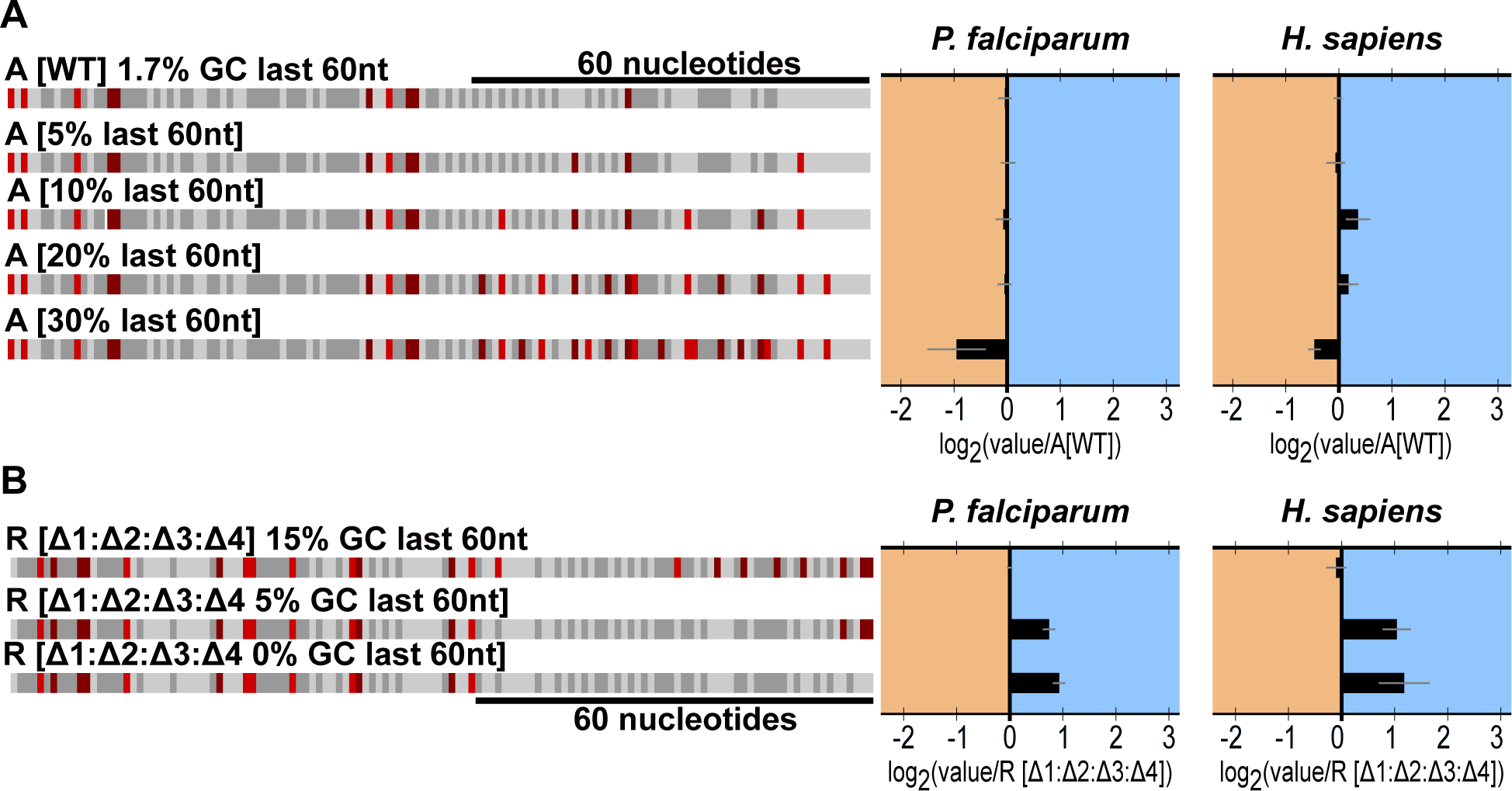
Dissecting the effects of the four uAUGs in R[WT]. uAUGs were found to have a generally repressive effect on TE that can be dependent on the presence each other. Graphed for each is the average and SEM of log_2_(each experimental value normalized to experimental R[WT] mean). The dotted line marks the average log_2_(R[Δ1Δ2Δ3Δ4] normalized to R[WT]). For each figure, to the left is a diagram of the sequences using the same annotations as Figure 2C. To the right of the diagrams are the results for *P. falciparum* and human K562s. a) The effect of removing all four uAUGs from R[WT]. b) The effect of retaining a single uAUG c) The effect of removing each uAUG individually d) The effect of removing two uAUGs in combination

To further evaluate the repressive effects of uAUGs in a novel context, the four uAUGs from R[WT] were placed into A[WT] at the matching positions (Supplemental Figure 3). As expected, in *P. falciparum*, when all four uAUGs were present A[+1:+2:+3:+4], translation was repressed, 2.9-fold. Additionally, each uAUG individually repressed translation between 1.5-fold and 2.9-fold when the other positions were mutated to “AUC (Supplemental Figure 3). The results in K562 followed the same trends as *P. falciparum*.

To explore potential interactions between uAUGs, pairwise combinations of the uAUGs in R[WT] were evaluated (Figure 3C). If uAUGs possess independent repressive potentials that do not affect each other, the repression by any two uAUGs would be the product of their respective potentials. For example, the two furthest uAUGs, uAUG-1 and uAUG-4, yielded 37% and 73% of the maximum translation of the derepressed construct R[Δ1Δ2Δ3Δ4] in *P. falciparum* lysates. Thus, if acting independently, the predicted yield for a 5’ UTR containing both uAUGs would equal 0.37 * 0.73, or 27%, of the maximum signal. The measured signal for this combination (R[Δ2Δ3]) was extremely close to the predicted value, 28.6%, suggesting that these two elements act independently and proportionately on translation. Evaluation of the remaining pairs of uAUGs revealed some notable combinations that likely highlight interacting pairs (Supplemental Figure 4). Of note, the predicted combination of uAUG-3 and uAUG-4 (R[Δ1Δ2]) in *P. falciparum* underestimates the measured amount of translation (11% predicted versus 19% measured), suggesting an interaction between uAUG-4 and uAUG-3, which, as noted previously, marks the end of the uORF formed by uAUG-4. For K562 lysates, constructs containing uAUG-2 differ most from their predicted values, indicating this element may be uniquely sensitive to the presence of the other uAUGs.

Having examined all pair-wise combinations of the four uAUGs, each three-way combination was then evaluated (Figure 3D). Unlike the broad range of differing repressive activities observed for individual and pairwise uAUGs, trios of uAUGs all repressed translation to a similar or greater degree than R[WT]. Together these data indicated that uAUGs in isolation independently confer varying levels of repression; however, multiple uAUGs may combine to produce a concerted effect that was not predicted by their individual contributions.

### Investigating the effect of position and termination status on uAUG repression

Each of the uAUGs in R[WT] is distinct with respect to their Kozak context, their position relative to the translation start site, and their termination status. Previous work describing the Kozak context for *P. falciparum* suggests a string of adenosine bases preceding the start site is most commonly observed [28, 33]. To assess the effects of uAUG positionality while maintaining a common Kozak, a cassette comprised of the -3 to +9 sequence from uAUG-3 was individually placed at five equally spaced positions within R[Δ1Δ2Δ3Δ4] beginning at -14 nucleotides from the reporter protein coding region (Figure 4). All cassettes were inserted in the +2 frame such that if translation initiated at these sites, no reporter should be translated in-frame. Two versions of the cassette were created, one maintaining the termination with a stop codon at the end of the cassette and one without (Figure 4A/B). For the five constructs containing a non-terminating uAUG, all potential stop sites proceeding the protein coding region in-frame with the 5’ most cassette were eliminated and the effect of these mutations alone in the presence of uAUG-3 (R[Δ1Δ2Δ4]*) were evaluated (Supplemental Figure 5A).

**Figure 4:**
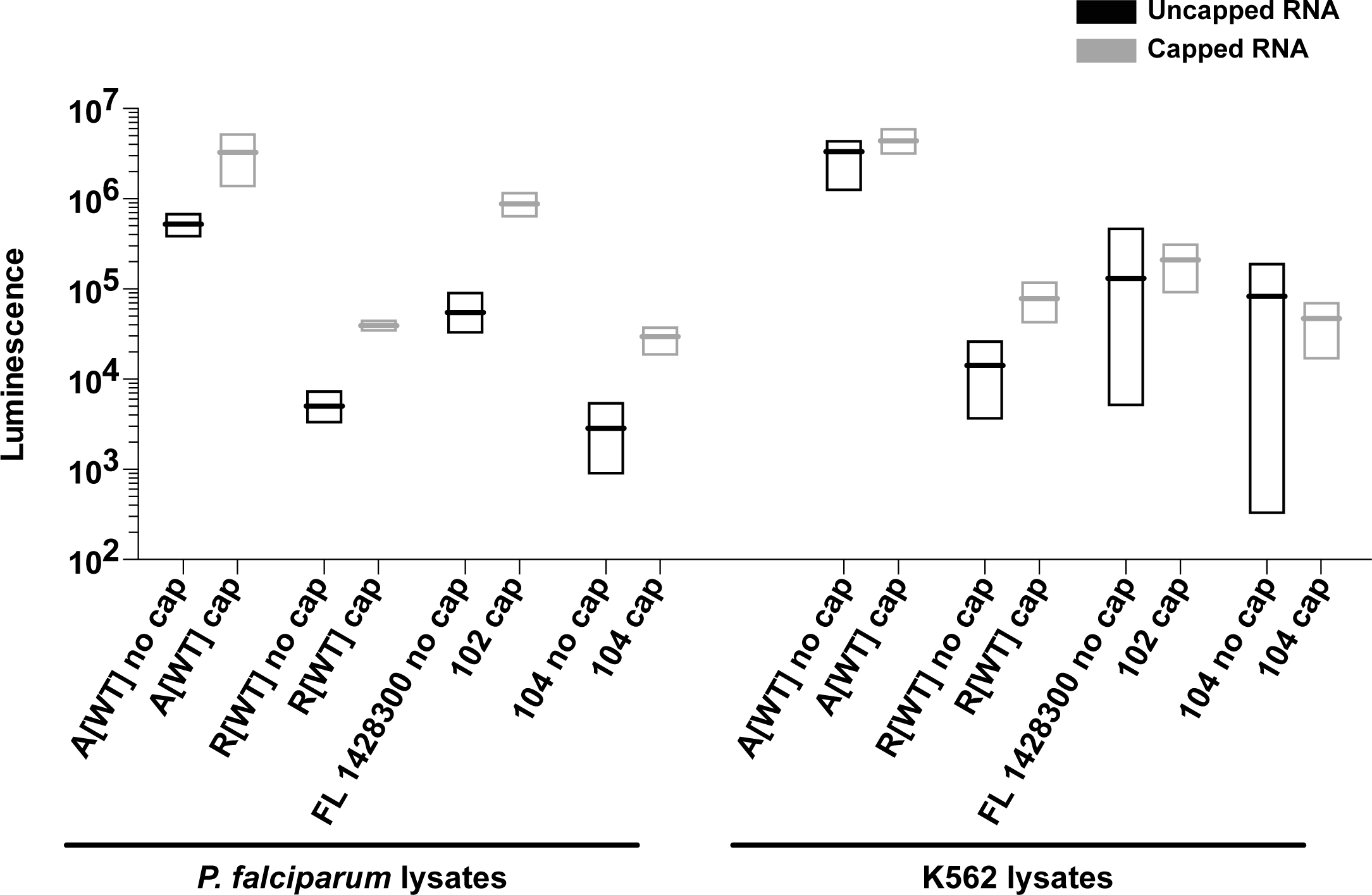
Effect of equally spaced and out of frame, non-terminated uAUGs or uORFs on TE a) Sequence diagram of the two cassettes inserted into R[Δ1:Δ2:Δ3:Δ4] at 5 different positions. The green arrows mark the uAUGs, the solid line indicates the length of the uORF, and the dotted line marks the sequence downstream of the non-terminated uAUG b) The sequence diagrams to the left represent the 5’ UTRs containing the uORF cassette. To the right the uORF cassette and the non-terminated cassette are presented side by side. The left set is from *P. falciparum* lysates while the right is from human. Graphed for each is the average and SEM of log_2_(each experimental value/ experimental mean of R[Δ1:Δ2:Δ3:Δ4]).

Except for the -122 position, where the uAUG is 11 nucleotides from the 5’ cap, all cassette placements resulted in repression comparable to R[Δ1Δ2Δ4] (Figure 4C). Of note, the cassettes placed nearest to the 5’ cap had little effect on translation in either *P. falciparum* or K562 lysates (1.2-fold and 1.3-fold repression respectively). For *P. falciparum*, unlike the relative consistency of repression produced by uORF placement, the uAUG equivalent yielded a trend in repression. As the uAUG moved closer to the translation start site the repressive strength increased until maximum repression was achieved when the cassette was placed -41 nucleotides from the translation start site (Figure 4C). In comparison, K562 lysates also yielded peak repression at the -41 position, but the pattern of repression induced by both the uORF and uAUG cassettes were more similar to each other and the trend observed for uAUG cassettes in *P. falciparum*. These experiments indicate that in both *P. falciparum* and K562 lysates, the position of uAUGs contributes in part to downstream repression, however, termination status may also impact this effect, at least in the case of *P. falciparum*.

### Evaluating the effect of GC content on TE

One distinguishing feature of the *P. falciparum* genome is an extreme bias in nucleotide content, especially within the intergenic regions that are ∼90% AT [34]. As noted in Figure 1D and 1E, there is a significant difference in the distributions of GC content between the 5’ UTRs of genes with high and low TE with repressed genes exhibiting a higher GC bias. These differences are evident within A[WT] and R[WT], which possess 7.7% GC, and 16.9% GC respectively. This GC bias is intensified in the 60 nucleotides closest to the translation start with A[WT] containing only 1.7% GC and R[WT] containing 15% GC (Figure 2D). To investigate the impact of GC content in the context of these two constructs, substitutions were systematically introduced into the proximal region of A[WT] to increase the GC content from 1.7% to a maximum of 30% GC (Figure 5A). Substitutions were maintained between constructs, no upstream “AUG”s were introduced, and significant secondary structures was avoided (Supplementary Figure 5). In *P. falciparum* lysates, between 1.7% and 20% GC there was no change in TE while at 30% GC translation was repressed 1.5-fold (Figure 5A). The repressive effect of the high GC content was 1.3-fold in human K562 lysates.

**Figure 5:**
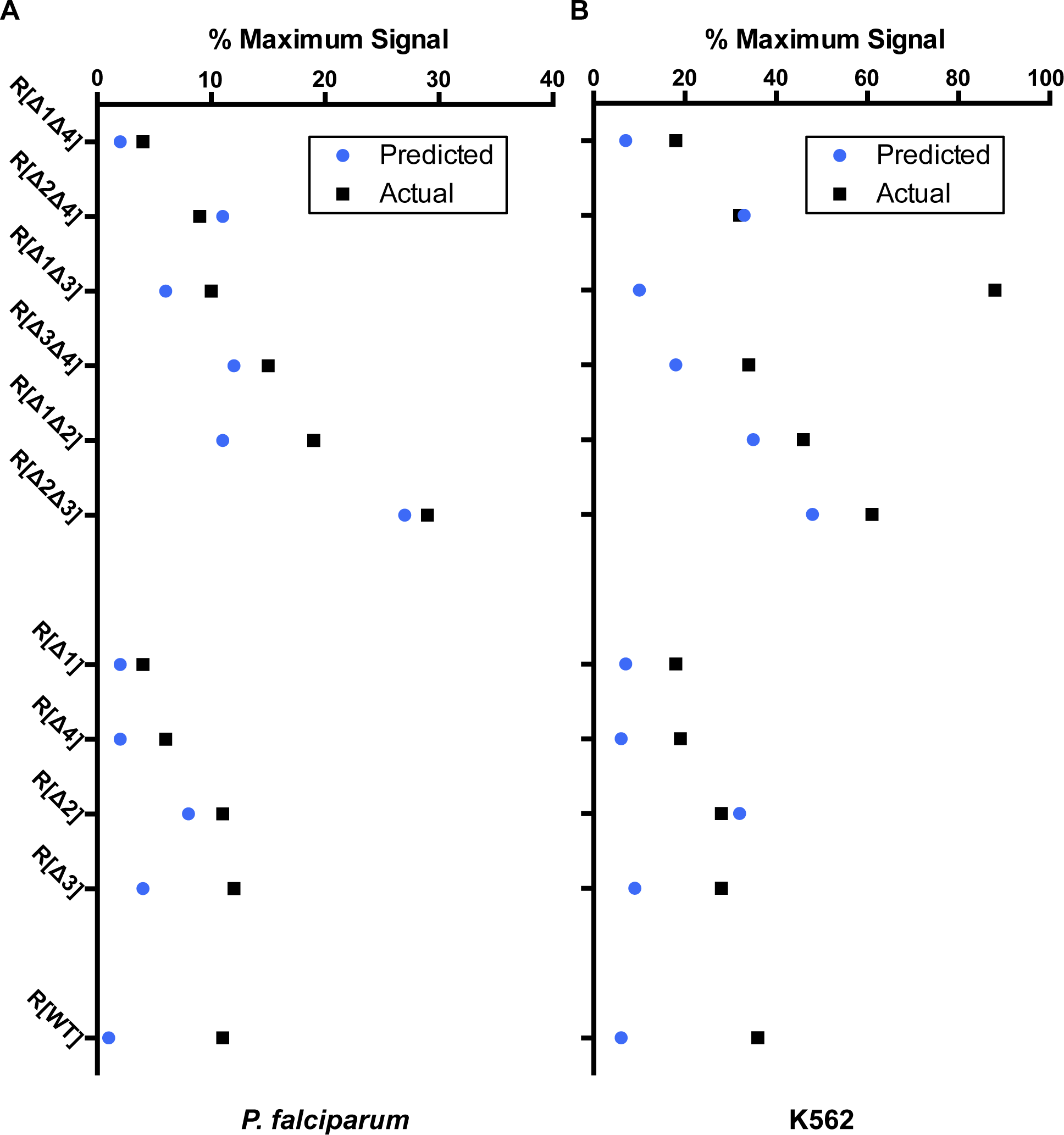
Evaluating the effect of GC content on translation a) Increasing GC content in A[WT]. Graphed for each is the average and SEM of log_2_(each experimental value/ experimental mean of A[WT]) b) eliminating GC content in R[Δ1:Δ2:Δ3:Δ4]. Graphed for each is the average and SEM of log_2_(each experimental value/ experimental mean of R[Δ1:Δ2:Δ3:Δ4])

The converse experiment of reducing the GC content of R[WT] was also carried out. The GC content in the last 60 nucleotides of R[Δ1Δ2Δ3Δ4] was reduced to 5% by eliminating all GC content between 4 and 60 nucleotides from the translation start site and to 0% by removing all GC (Figure 5B). A maximum translation increase of approximately 2-fold was observed relative to R[Δ1Δ2Δ3Δ4], indicating a modest but measurable impact in this context. These results were mirrored in K562 lysates (Figure 5B). Together, the result of manipulating the GC content of the last 60 nucleotides of the 5’ UTR suggests that the impact on translation to be subtle, but sensitive to the overall context.

### Identifying additional *cis*-acting regulatory regions within R[WT] and A[WT]

In addition to the study of specific elements predicted to impact TE, a series of systematic sequence swaps were investigated, in which regions from both the 5’ and 3’ end of R[WT] and A[WT] were exchanged. Beginning with the 3’ end of the 5’ UTR, 20, 40, and 60 nucleotides were exchanged between R[WT] and A[WT] (Figure 6A and 6B). In the case of A[WT], introducing more sequence from R[WT] severely impacted TE. While some of this impact was anticipated due to the introduction of uAUG-1 and uAUG-2, additional decreases in translation were observed with sequence beyond these elements (A[60nt 3’ R]). Furthermore, the added impact beyond the introduction of uAUGs was observed only with *P. falciparum* lysates. For the converse experiments, exchange of sequence from A[WT] into R[WT] at the 3’ end resulted in increased translation (11.7-fold). This increase in translation was in part expected due to the elimination of uAUG-1 and uAUG-2, however the magnitude of the effect is greater than predicted from the experiments shown in Figure 3C. The effect in human K562 lysates was markedly less with a maximum difference of 1.4-fold.

**Figure 6:**
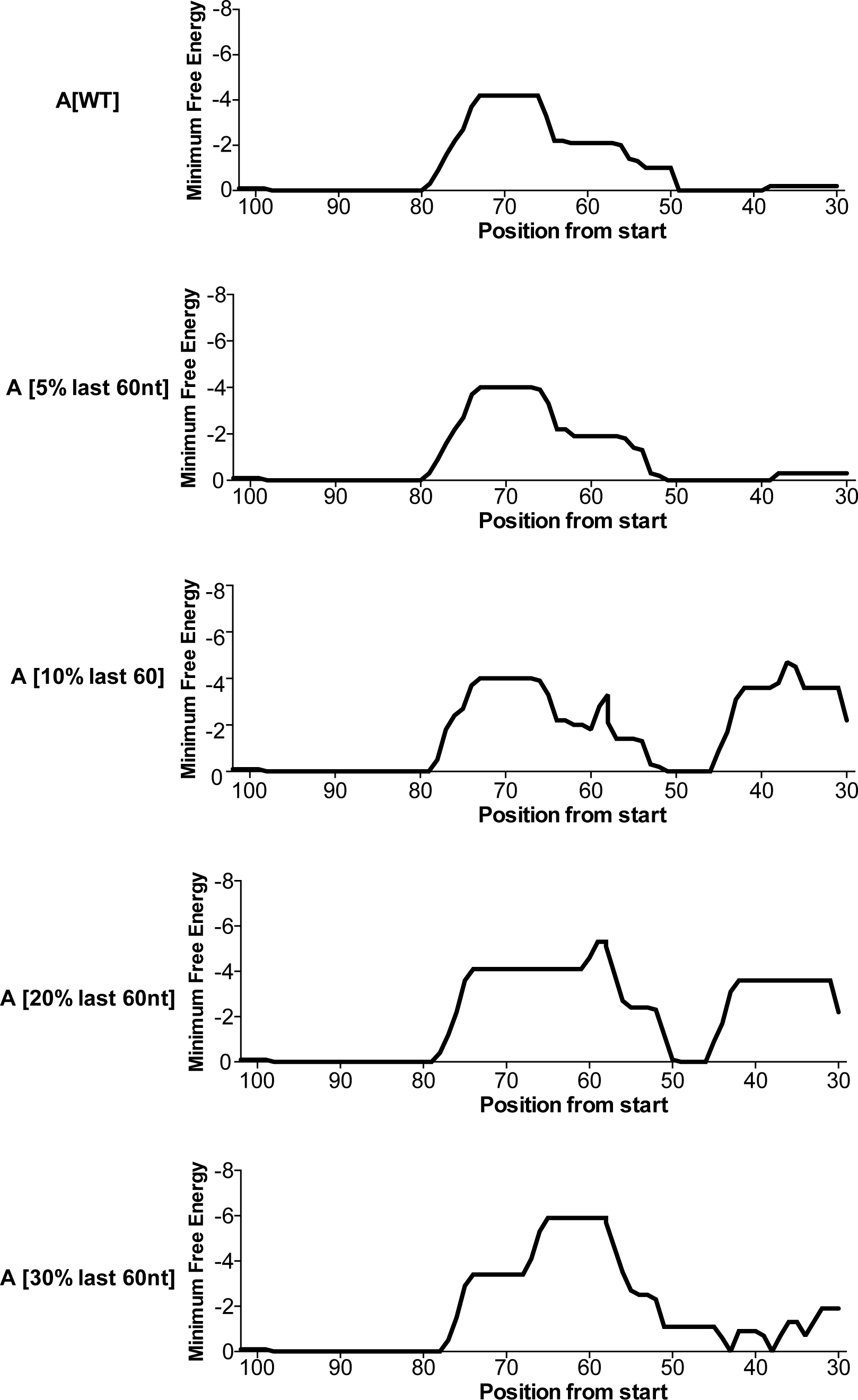
Evaluating the effects of the ends of the 5’ UTRs on translation a) Swapping the 3’ end of R[WT] into A[WT] Graphed for each is the average and SEM of log_2_(each experimental value/ mean experimental A[WT]) b) swapping the 3’ end of A[WT] into R[WT]. Graphed for each is the average and SEM of log_2_(each experimental value / mean experimental R[WT]) c) swapping the 5’ end of R[Δ1:Δ2:Δ3:Δ4] into A[WT]. Graphed for each is the average and SEM of log_2_(each experimental value / mean experimental of A[WT]) d) swapping the 5’ end of A[WT] into R[WT]. Graphed for each is the average and SEM of log_2_(each experimental value / mean experimental R[Δ1:Δ2:Δ3:Δ4]).

Sequence exchanges at the 5’ end were similarly carried out using 10, 20, and 30 nucleotide swaps between A[WT] and R[Δ1:Δ2:Δ3:Δ4]. The latter construct was chosen over R[WT] to assess the impact in the absence of uAUGs. For *P. falciparum*, exchanging the first 10 nucleotides of R[Δ1:Δ2:Δ3:Δ4] into A[WT] repressed translation 2.6-fold, with a final 3.7-fold repression exchanging 30 nucleotides. (Figure 6C). In contrast, exchanging the first 10 and 20 nucleotides of A[WT] into R[Δ1:Δ2:Δ3:Δ4] activated translation up to 1.9-fold while exchanging 30 nucleotides activated translation 3.5-fold. Note that the level of translation achieved in this latter construct matches the output of A[WT], demonstrating that in the absence of uAUGs, exchanging the sequence elements within the first 30 nucleotides of the 5’ end of the 5’ UTR was sufficient to render A[WT] and R[Δ1:Δ2:Δ3:Δ4] approximately equivalent (Figure 6D).

## Discussion

Among eukaryotes, *P. falciparum* presents several distinct features that bear upon translation. First, the AT-rich genome contains frequent poly-adenosine stretches that alone necessitates unique adaptions of the translational machinery to prevent ribosome stalling or frameshifting [35, 36]. Additionally, there are a limited number of ribosomal RNA copies within the genome, each with stage specific expression [37, 38]. The transcriptome also features unusually long 5’ UTRs, the longest in late trophozoites being a remarkable 8229 nucleotides (PF3D7_1139300). Despite these features, previous studies suggest that *P. falciparum* initiates translation in a cap-dependent manner similarly to other eukaryotes [39, 40].

While the central initiation factors required for cap-binding have been bioinformatically identified and many of the essential interactions have been validated, questions remain around how these factors regulate translation initiation given *P. falciparum*’s unique 5’ UTR features [26, 41]. Additionally, ribosome profiling has demonstrated that translation is an integral point of regulation for model eukaryotes [42, 43], but for *P. falciparum* it reveals that less than 10% of transcripts are translationally regulated. Directly evaluating how these unusual mRNA features function in *P. falciparum* could reveal unique mechanisms that would be powerful therapeutic targets.

A re-analysis of ribosome profiling data highlights two important features that differ between mRNAs at the top and bottom of the TE range. As shown in Figure 1, the presence of uAUGs and GC content are significantly different between highly translated and poorly translated mRNAs, a difference that appears exacerbated by proximity to the protein coding region. To explore and dissect the role of these features, two representative 5’ UTRs were chosen from the top and bottom deciles, the 5’ UTRs of PF3D7_1411400 and PF3D7_1428300. The differences in TE driven by these two 5’ UTRs were faithfully recapitulated using *in vitro* translation extracts generated from late trophozoites of *P. falciparum* W2 strain (Figure 2) and human K562 cells. Surprisingly, these differences were maintained when using only the proximal 130 nucleotides from each 5’ UTR, with A[WT] derived from PF3D7_1428300 and R[WT] from PF3D7_1411400. These two 130-nucleotide 5’ UTRs provided an ideal platform to evaluate the effects of uAUGs and GC content.

uAUGs have long been appreciated as translational regulatory elements, and work by Marilyn Kozak demonstrated their repressive abilities in the early 1980s [20]. However, it remains difficult to predict the individual or joint repressive activities of uAUGs from sequence context alone, especially for non-model organisms. Additionally, it is unusually to have uAUGs so abundant throughout the transcriptome. Here, a reductionist approach was used to individually assess the repressive potential of each uAUG within R[WT] in isolation, and in combination (Figure 3). For many pairs, such as uAUG-1 and uAUG-4, the combined activity directly reflected a combination of each uAUG’s repressive strength. For others, like uAUG-3 and uAUG-4, it was revealed that the combined effect of two uAUGs could be reduced by their interaction. Since uAUG-3 is itself the in-frame stop site for uAUG-4, it reasonable to assume that the termination of uAUG-4 may interfere with initiation events at uAUG-3. These interactions make it difficult to predict the impact of multiple uAUGs without direct measurements as performed here.

The sequence context surround an “AUG” is essential for determining the rate of initiation at that site [11,44,45], however, additional elements may affect the regulatory activity of an uAUG. Here, two possible modifiers were examined in detail, namely, the position of uAUGs relative to the protein coding region, and whether it forms a uORF (Figure 4). Both the position and termination status affect translation with the most dramatic result arising when the uAUG is positioned furthest from the protein coding region, only 11 nucleotides from the 5’ cap. At this distance neither the open uAUG nor the uORF repressed translation. One caveat of this study is that only one putative uORF was assessed. It is likely that the length and composition of the uORF sequence itself may modify the overall impact.

Along with uAUG frequency, bioinformatic analysis of the 5’ UTR sequences from *P. falciparum* also reveals a statistically significant difference in GC content, with higher GC content corresponding to lower TE. While higher GC content could correlate with higher secondary structures, we wanted to evaluate if GC content alone could regulate translation. Surprisingly, the results of manipulating GC content proximal to the protein coding region in the context of only these two chosen UTRs yielded corresponding changes in the predicted direction, albeit with small magnitudes when compared to the impact of uAUGs. In the active context, translation became repressed relative to A[WT] at 30% GC content within 60 nucleotides of the translational start. Within this 60-nucleotide region, only 31 (1.4%) of the 2088 5’ UTRs from *P. falciparum* expressed in late trophozoites evaluated here are 30% GC or above (Figure 1A). Thus, few genes would be predicted to be impacted by these shifts in GC content alone. Eliminating GC content from the last 60 nucleotides of R[Δ1:Δ2:Δ3:Δ4] resulted in modest increases in TE (Figure 5B). In this case, of the 2088 5’ UTRs 273 (13.1%) are 5% or below within this region and 16 (0.8%) are 0%.

Finally, to examine the effects of the sequences within A[WT] and R[WT]/ R[Δ1:Δ2:Δ3:Δ4] on translation, segments from the 5’ and 3’ ends were progressively exchanged between them (Figure 6). Sequence exchanges at the 3’ end of the 5’ UTR removed or introduced uAUGs, which resulted in the expected increases or decreases in TE respectively. We note that in each case, exchanged sequence beyond the uAUGs also impacted TE in *P. falciparum*, suggesting additional context within these regions. Sequence exchanges at the 5’ end were more impactful than would have been predicted. Specifically, 30 nucleotides of A[WT], when substituted into R[Δ1:Δ2:Δ3:Δ4], suggest a possible sequence with a role in regulating the rate of translation initiation.

As an essential pathway throughout the parasite’s life cycle, protein synthesis is an attractive therapeutic target. However, since the mechanisms of eukaryotic translation are highly conserved, potential therapeutics must cross the challenging bar of being highly specific to *P. falciparum*. Here, *in vitro* translation was used to allow for direct comparison between *P. falciparum* and human to identify unique effects on TE. Despite the large evolutionary distance between the two organisms, *P. falciparum* and K562 lysates yielded highly similar results in the context of the two short model UTRs used here. For developing therapeutics targeting translation initiation, avoiding host effects will be challenging, but *in vitro* translation can continue to be a valuable tool to directly measure differences between Plasmodium and humans [46].

Finally, this work continues the task of uncovering the complexity of 5’ UTR *cis*-acting regulatory elements and their impact on TE in eukaryotes. *In vitro* translation has previously revealed the importance of the Kozak consensus sequence and uAUGs in model eukaryotes, such as Saccharomyces cerevisiae and mammalian cultures [21,47–49], while higher throughput selection and machine learning techniques have been used to probe the effect of 5’ UTR *cis*-acting elements in these same systems [24, 50]. However, working with non-model organisms such as *P. falciparum* poses unique challenges, such that many of these techniques cannot be readily utilized for comparative analysis. The highly reductionist approach taken here has the benefit of allowing specific and systematic hypotheses to be tested, although it is clear that higher throughput methods will be required to generalize these findings beyond these specific examples.

## Conclusions

*Cis*-acting features within the 5’ UTRs of eukaryotes regulate the TE of a given gene. While specific examples have previously been evaluated in model eukaryotes, *P. falciparum* possesses unusual 5’ UTR characteristics, such as length, base content, and high uAUG prevelence, that suggest *cis*-acting upstream elements play a significant role in tuning translational efficiencies. Through extensive dissection of exemplar 5’ UTRs from *P. falciparum*, we measure the individual impacts of each putative element while comparing these same constructs in human lysates. The impact of these elements was found to be surprisingly similar in both systems. Since, unlike humans and most other studies eukaryotes, long 5’ UTRs featuring multitudes of uAUGs are common in *P. falciparum*, the precise configuration of these elements may have evolved to tune translation levels in this organism where other post-transcriptional regulatory mechanisms may be absent.

## List of abbreviations

uAUG: upstream “AUG”
uORF: upstream open reading frame
5’ UTR: 5’ untranslated region
TE: translational efficiency

## Declarations

Ethics approval and consent to participate

Consent for publication

## Availability of data and material

The datasets supporting the conclusions of this article are included within the article in Supplemental File 4. The TEs and 5’ UTR sequences from that data used for comparative analysis here can be found in Supplemental File 1. The previously published data from Caro, Ahyong *et. al.* [18] can be found at available at Dryad Digital Repository under a CC0 Public Domain Dedication: http://dx.doi.org/10.5061/dryad.vb855.

## Authors’ Contributions

VEG and JLD conceived and designed this study. VEG performed and executed the experiments. RD maintained, harvested, and generated *in vitro* translation lysates for the K562 cells. VEG and JLD drafted and edited this manuscript. All authors read and approved the submitted manuscript.

## Acknowledgements

We would like to acknowledge the DeRisi lab’s Team Malaria for advice, thoughts, and training. Additionally, we would like to thank Yun Song and Adam Frost for valuable discussion and comments on the manuscript and Hanna Retallack, Jamin Lui, Madhura Raghavan, Sara Sunshine, Elze Rackaityte, and Caleigh Mandle-Brehm for their edits and commentary on the paper.

## Figure Legends

**Supplemental Figure 1:**
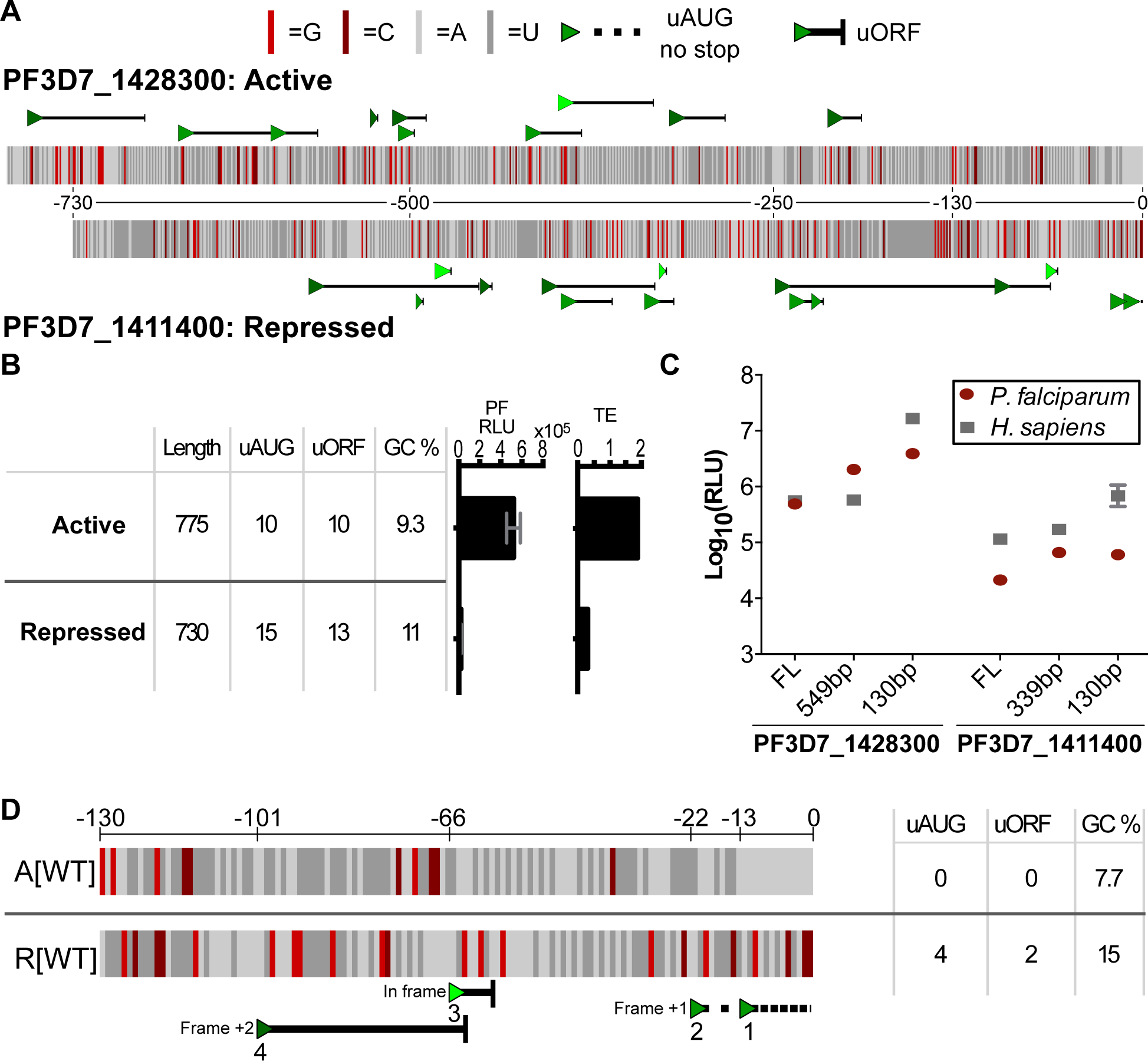
Further comparison of 5’ UTR features of genes in the bottom 10% and top 10% of TEs in the late trophozoite stage using data from Caro and Ahyong *et. al.* 2014 [18]. a) Distributions of the 5’ UTR lengths from genes with high (blue) or low (yellow) TE. KS test. statistic comparison of the two: 0.12 p-value 0.1 b) Distributions of the total number of uAUGs in the 5’ UTRs from genes with high (blue) or low (yellow) TE. KS test statistic comparison of the two: 0.31 p-value 1.5*10^-5^.

**Supplemental Figure 2:**
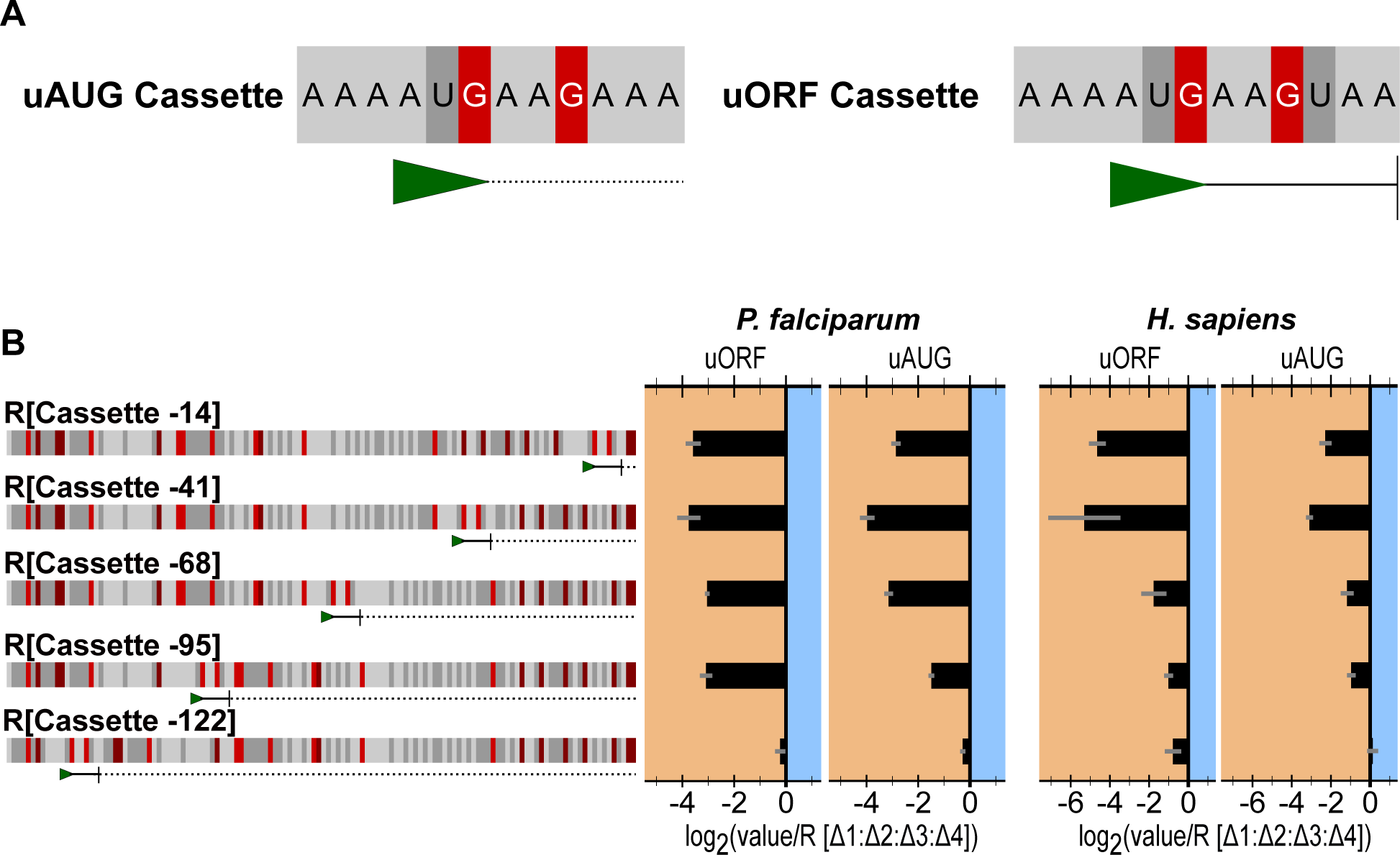
The raw luminescence signal from capped and uncapped RNAs in *P. falciparum* and K562 *in vitro* translation.

**Supplemental Figure 3:**
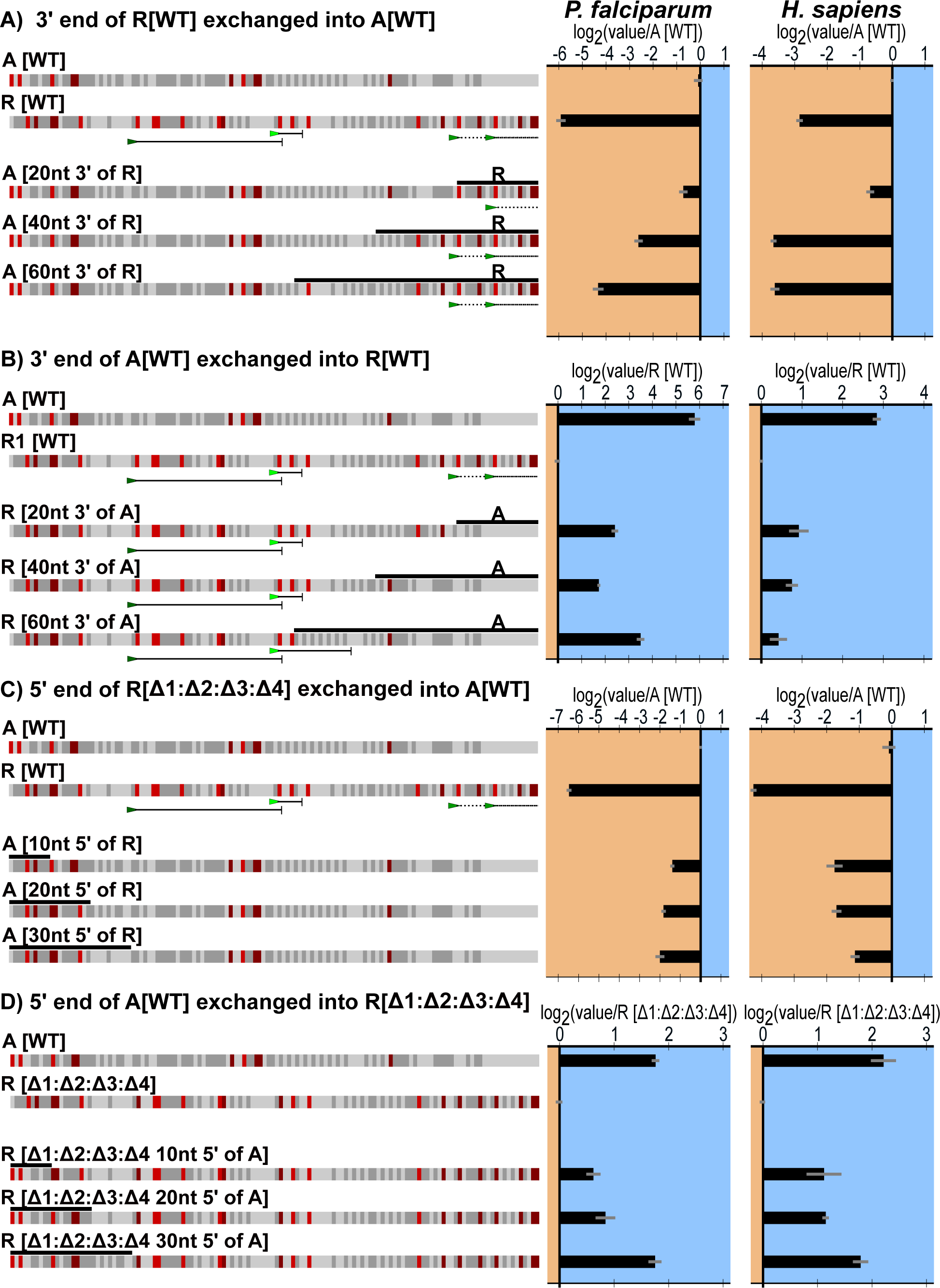
The 4 uAUGs from R[WT] exchanged into A[WT] at the same positions showing that the repressive effect is conferrable to other contexts. Graphed for each is the average and SEM of log_2_(each experimental value normalized to the experimental average of A[WT]).

**Supplemental Figure 4:**
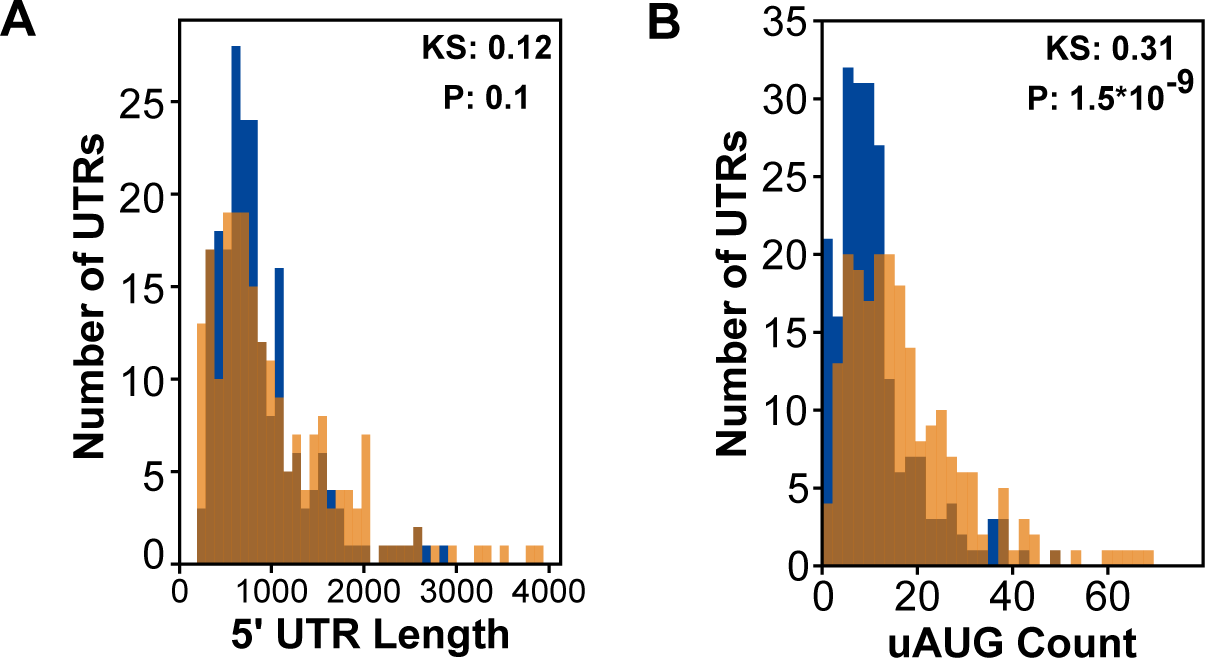
Predicted repressive effect of combinations of the uAUGs in R[WT] based on their individual activities for a) *P. falciparum* and b) K562.

**Supplemental figure 5:**
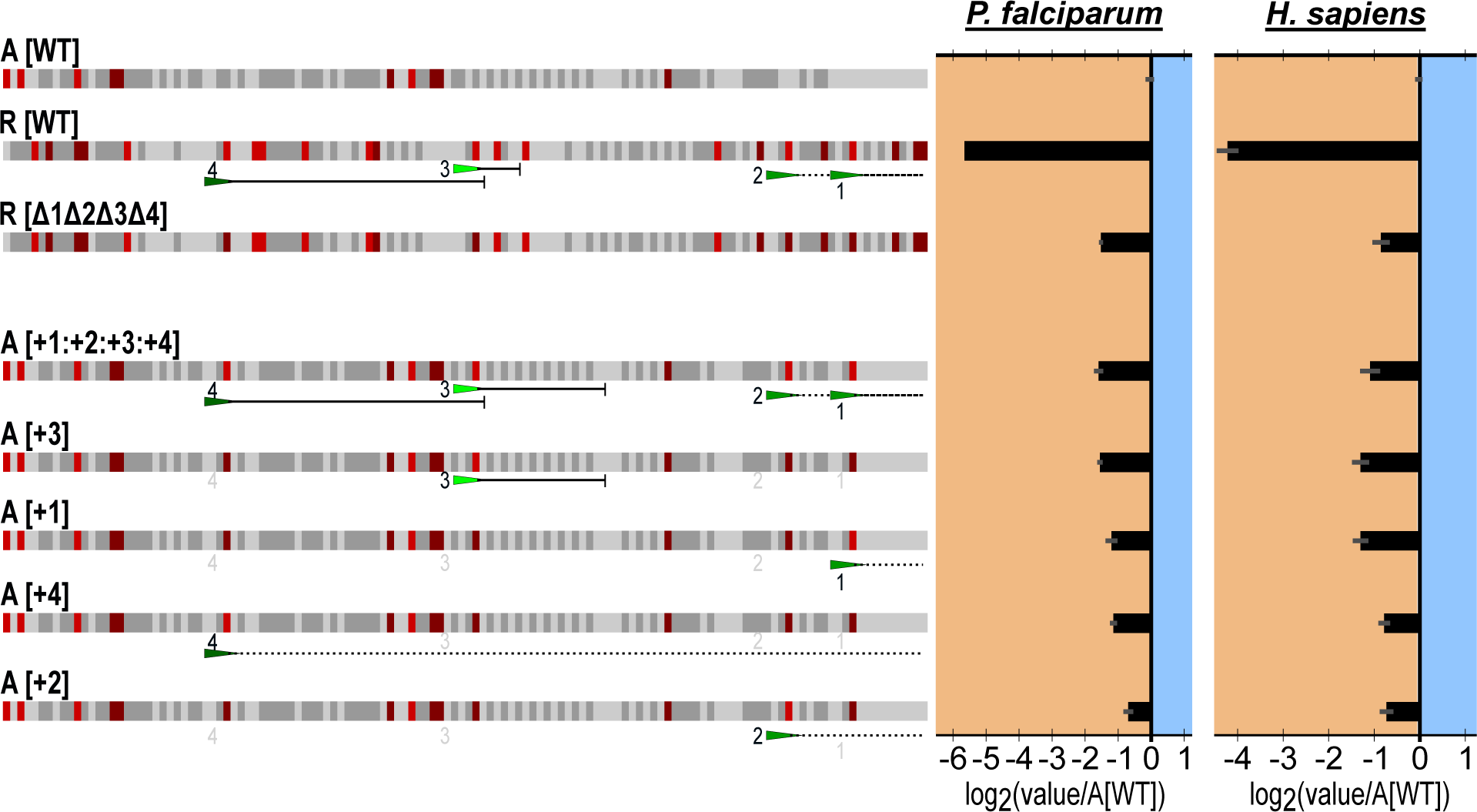
To eliminate all downstream stop sites for moving the out of frame non-terminated uAUG, 6-point mutations had to be added to the 5’ UTR. R[Δ1:Δ2:Δ4]^*^ was made with those point mutations to compare to R[Δ1:Δ2:Δ4]. Graphed for each is the average and SEM of log_2_(each triplicate value/ average of R[Δ1:Δ2:Δ3:Δ4] experimental triplicates)

**Supplemental figure 6:**
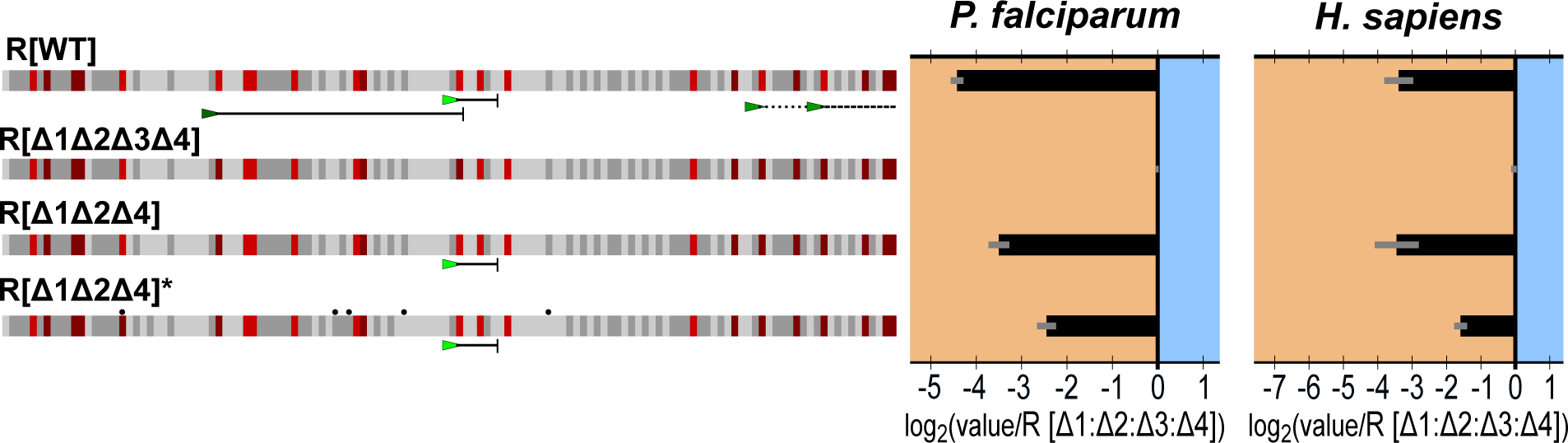
The predicted free energy of the secondary structure with in a 30-nucleotide sliding window moved by 1 nucleotide across the 5’ UTRs used to evaluate the effect of GC content.

## Additional Files

Additional File 1: 5’ UTR analyzed. The data on the 2088 genes used for the analysis in Figure 1 and Supplemental Figure 1, including the 5’ UTR sequences used, .xls.

Additional File 2: P16 sequence. The plasmid sequence for P16 used to generate new constructs, .geneious.

Additional File 3: 5’ UTR sequences. The sequences of all the 5’ UTR sequences evaluated in this manuscript, .fasta.

Additional File 4: Figure data. All the raw and processed data used to generate the figures in this manuscript, .xls.

Additional File 5: Figure generating script. The script used to generate the figures within the manuscript, .py.

