## Additional File 5 for "Functional characterization of 5’ UTR *cis*-acting sequence elements that modulate translational efficiency in *P. falciparum* and humans": Additional_File_5.py

```
#! /usr/bin/env python
# python seqplot-v4_takesFASTA, -f fasta file
#currently creates one file per sequence

#This code takes in three files, a fasta file containing the sequences needing to be graphed (-f), the plamsodium data in a csv with the columns labeled with the gene names and the results below (-p), and the human data in a csv with the columns labeled with the gene names and the results below (-h). An output file name is needed (-o) that needs to end in .eps since an eps file will be generated.

import sys, getopt, pandas

from psfile import EPSFile

genename=""
sequence=""
boxwidth = 5

#Code to create the AUG triangels
def startcodon(x,yposition,shade,fd,boxwidth):
    startx = x
    starty = yposition-10
    start_width=boxwidth*3
    fd.append("%f %f moveto"%(startx,starty))
    fd.append("[ ] 0 setdash")
    fd.append("0 7 rlineto")
    fd.append("%f -3.5 rlineto"%(start_width))
    fd.append("closepath")
    fd.append("gsave")
    fd.append("0 %f 0 setrgbcolor"%(shade))
    fd.append("fill")
    fd.append("grestore")
    fd.append("0 0 0 setrgbcolor")
    fd.append("0.5 setlinewidth")
    fd.append("stroke")
    return

#Code to create the uORF lines
def uORF(x,yposition,z,fd,boxwidth):
    startx = x +boxwidth*2
    starty = yposition-6.5
    uORF_length=z
    fd.append("3 setlinewidth")
    fd.append("%f %f moveto"%(startx,starty))
    fd.append("%f 0 rlineto"%(uORF_length))
    fd.append("stroke")
    startx = startx+z
    starty = yposition-12
    fd.append("2 setlinewidth")
    fd.append("%f %f moveto"%(startx,starty))
    fd.append("0 10 rlineto")
    fd.append("stroke")
    return


#Code adds dotted line after AUGs that don't have a stop
def nostop(x,yposition,fd,seqlength, distance_from_end,boxwidth):
    startx = x +boxwidth*2
    starty = yposition-6.5
    length_line=distance_from_end*boxwidth-boxwidth*2
    fd.append("3 setlinewidth")
    fd.append("[3 5 3 5] 0 setdash")
    fd.append("%f %f moveto"%(startx,starty))
    fd.append("%f 0 rlineto"%(length_line))
    fd.append("stroke")
    fd.append("stroke")
    return


#Code to add the UTR name above the image
def addname(x,y,genename):
    starty=y+boxheight+5
    fd.append("0 setgray")
    fd.append("/Helvetica 20 selectfont")
    fd.append("%f %f moveto"%(x, starty))
    fd.append("(%s) show"%genename)
    return

#Code do draw the data bar for the graph of plasmodium data
def pf_add_data_bar(pf_mean_translation, pf_sem_translation, y, pf_y_axis_x, pf_tick_mark_distance):
    y_axis_x=pf_y_axis_x
    bar_length=pf_mean_translation*pf_tick_mark_distance
    neg_bar_length=0-bar_length
    #The code below draws the bar of the bar graph
    fd.append("%f %f moveto"%(y_axis_x,y))
    fd.append("[ ] 0 setdash")
    fd.append("%f 0 rlineto"%(bar_length))
    fd.append("0 15 rlineto")
    fd.append("%f 0 rlineto"%(neg_bar_length))
    fd.append("closepath")
    fd.append("gsave")
    fd.append("0 setgray")
    fd.append("fill")
    fd.append("grestore")
    fd.append("0 0 0 setrgbcolor")
    fd.append("0.5 setlinewidth")
    fd.append("stroke")
    
    #The code below adds the SEM bars
    sem_start_x=y_axis_x+bar_length
    sem_start_y=y+7.5
    sem_x_length=pf_sem_translation*pf_tick_mark_distance
    fd.append("%f %f moveto"%(sem_start_x,sem_start_y))
    fd.append("0.5 setgray")
    fd.append("2.5 setlinewidth")
    #fd.append("0 setgray")
    fd.append("%f 0 rlineto"%(sem_x_length))
    fd.append("%f %f moveto"%(sem_start_x,sem_start_y))
    fd.append("%f 0 rlineto"%(-sem_x_length))
    fd.append("stroke")
    return

#Code do draw the data bar for the graph of human data
def human_add_data_bar(human_mean_translation, human_translation, y, human_y_axis_x, human_tick_mark_distance):
    y_axis_x=human_y_axis_x
    bar_length=human_mean_translation*human_tick_mark_distance
    neg_bar_length=0-bar_length
    #The code below draws the bar of the bar graph
    fd.append("%f %f moveto"%(y_axis_x,y))
    fd.append("[ ] 0 setdash")
    fd.append("%f 0 rlineto"%(bar_length))
    fd.append("0 15 rlineto")
    fd.append("%f 0 rlineto"%(neg_bar_length))
    fd.append("closepath")
    fd.append("gsave")
    fd.append("0 setgray")
    fd.append("fill")
    fd.append("grestore")
    fd.append("0 0 0 setrgbcolor")
    fd.append("0.5 setlinewidth")
    fd.append("stroke")
    
    #The code below adds the SEM bars
    sem_start_x=y_axis_x+bar_length
    sem_start_y=y+7.5
    sem_x_length=human_sem_translation*human_tick_mark_distance
    fd.append("%f %f moveto"%(sem_start_x,sem_start_y))
    fd.append("0.5 setgray")
    fd.append("2.5 setlinewidth")
    #fd.append("0 setgray")
    fd.append("%f 0 rlineto"%(sem_x_length))
    fd.append("%f %f moveto"%(sem_start_x,sem_start_y))
    fd.append("%f 0 rlineto"%(-sem_x_length))
    fd.append("stroke")
    return

def make_axis(pf_min_x, pf_max_x, human_min_x, human_max_x, count, end_of_sequence_diagrams, canvaslength, pf_y_axis_x, pf_tick_mark_distance, human_y_axis_x, y_axis_y, human_tick_mark_distance):
    #WE WILL START WITH THE PLASMODIUM/LEFT DATA SET
    number_of_tick_marks=round(200/pf_tick_mark_distance)
    #This adds a first set of axis to the right of the sequence images but to the left of the second set
    #This adds y-axis to the right of the images
    print("count" + str(count))
    y_axis_length=(count)*27.5 #may want to play with this later *******************
    fd.append("5 setlinewidth")
    fd.append("%f %f moveto"%(pf_y_axis_x,y_axis_y))
    fd.append("0 %f rlineto"%(-y_axis_length))
    fd.append("stroke")

    #This adds the x-axis on the bottom
    x_axis_x=end_of_sequence_diagrams+5
    bottom_x_axis_y=canvaslength-y_axis_length-19 #The -30 accounts for how far down the y axis starts (in y_axis_y)
    fd.append("3 setlinewidth")
    fd.append("%f %f moveto"%(x_axis_x,bottom_x_axis_y))
    fd.append("225 0 rlineto")
    fd.append("stroke")

    
    #This adds tick marks to the bottom x-axis
    fd.append("1 setlinewidth")
    bottom_tick_mark_y=bottom_x_axis_y-5
    bottom_label_y=bottom_tick_mark_y-12
    tick_mark=int(pf_min_x)
    tick_mark_x=end_of_sequence_diagrams + 20
    bottom_label_x=tick_mark_x-9
    current_tick_mark=-1 #Can change this number to minipulate how many tick marks there are
    while current_tick_mark<number_of_tick_marks:
        fd.append("%f %f moveto"%(tick_mark_x,bottom_tick_mark_y))
        fd.append("0 10 rlineto")
        fd.append("stroke")
        fd.append("0 setgray")
        fd.append("/Helvetica 20 selectfont")
        fd.append("%f %f moveto"%(bottom_label_x,bottom_label_y))
        #fd.append("(%s) show"%(tick_mark))

        tick_mark+=1
        current_tick_mark+=1
        tick_mark_x+=pf_tick_mark_distance
        percent=str(tick_mark/2)
        bottom_label_x=tick_mark_x-4

    #This adds the x-axis on the top
    top_x_axis_y=canvaslength-30
    fd.append("5 setlinewidth")
    fd.append("%f %f moveto"%(x_axis_x,top_x_axis_y))
    fd.append("225 0 rlineto")
    fd.append("stroke")

    #This adds tick marks to the top x-axis
    fd.append("1 setlinewidth")
    top_tick_mark_y=top_x_axis_y-5
    top_label_y=top_tick_mark_y+12
    tick_mark=int(pf_min_x)
    tick_mark_x=end_of_sequence_diagrams + 20
    top_label_x=tick_mark_x-9
    current_tick_mark=-1
    while current_tick_mark<number_of_tick_marks:
        fd.append("%f %f moveto"%(tick_mark_x,top_tick_mark_y))
        fd.append("0 10 rlineto")
        fd.append("stroke")

        fd.append("0 setgray")
        fd.append("/Helvetica 20 selectfont")
        fd.append("%f %f moveto"%(top_label_x,top_label_y))
        fd.append("(%s) show"%(tick_mark))

        tick_mark+=1
        current_tick_mark+=1
        tick_mark_x+=pf_tick_mark_distance
        percent=str(tick_mark/2)
        top_label_x=tick_mark_x-4

    number_of_tick_marks=round(200/human_tick_mark_distance)
   
   #The code below creates a second set of axis to the right for the human data
   #NEXT WE HAVE THE HUMAN/RIGHT DATA SET
   #This adds y-axis to the right of the images
    print("count" + str(count))
    y_axis_length=(count)*27.5 #may want to play with this later *******************
    fd.append("5 setlinewidth")
    fd.append("%f %f moveto"%(human_y_axis_x,y_axis_y))
    fd.append("0 %f rlineto"%(-y_axis_length))
    fd.append("stroke")
    
    #This adds the x-axis on the bottom
    x_axis_x=end_of_sequence_diagrams+255
    bottom_x_axis_y=canvaslength-y_axis_length-19 #The -30 accounts for how far down the y axis starts (in y_axis_y)
    fd.append("3 setlinewidth")
    fd.append("%f %f moveto"%(x_axis_x,bottom_x_axis_y))
    fd.append("225 0 rlineto")
    fd.append("stroke")
    
    
    #This adds tick marks to the bottom x-axis
    fd.append("1 setlinewidth")
    bottom_tick_mark_y=bottom_x_axis_y-5
    bottom_label_y=bottom_tick_mark_y-12
    tick_mark=int(human_min_x)
    tick_mark_x=end_of_sequence_diagrams + 270
    bottom_label_x=tick_mark_x-9
    current_tick_mark=-1
    while current_tick_mark<number_of_tick_marks:
        fd.append("%f %f moveto"%(tick_mark_x,bottom_tick_mark_y))
        fd.append("0 10 rlineto")
        fd.append("stroke")
        fd.append("0 setgray")
        fd.append("/Helvetica 20 selectfont")
        fd.append("%f %f moveto"%(bottom_label_x,bottom_label_y))
        #fd.append("(%s) show"%(tick_mark))
        
        tick_mark+=1
        current_tick_mark+=1
        tick_mark_x+=human_tick_mark_distance
        percent=str(tick_mark/2)
        bottom_label_x=tick_mark_x-4
    
    #This adds the x-axis on the top
    top_x_axis_y=canvaslength-30
    fd.append("5 setlinewidth")
    fd.append("%f %f moveto"%(x_axis_x,top_x_axis_y))
    fd.append("225 0 rlineto")
    fd.append("stroke")
    
    #This adds tick marks to the top x-axis
    fd.append("1 setlinewidth")
    top_tick_mark_y=top_x_axis_y-5
    top_label_y=top_tick_mark_y+12
    tick_mark=int(human_min_x)
    tick_mark_x=end_of_sequence_diagrams + 270
    top_label_x=tick_mark_x-9
    current_tick_mark=-1
    while current_tick_mark<number_of_tick_marks:
        fd.append("%f %f moveto"%(tick_mark_x,top_tick_mark_y))
        fd.append("0 10 rlineto")
        fd.append("stroke")
        
        fd.append("0 setgray")
        fd.append("/Helvetica 20 selectfont")
        fd.append("%f %f moveto"%(top_label_x,top_label_y))
        fd.append("(%s) show"%(tick_mark))
        
        tick_mark+=1
        current_tick_mark+=1
        tick_mark_x+=human_tick_mark_distance
        percent=str(tick_mark/2)
        top_label_x=tick_mark_x-4
    
    return


#Get the command line options and opens the fasta file as a normal file, opens the data into a pandas data frame, and names the output
opts, args = getopt.getopt(sys.argv[1:],"f:p:h:d:o:")
for opt, arg in opts:
    if opt == "-f":
        myfile = arg
        #if opt =="-l":
        #fold_data = pandas.read_csv(arg)
    if opt== "-p":
        plasmo_data = pandas.read_csv(arg)
    if opt=="-h":
        human_data = pandas.read_csv(arg)
    if opt == "-o":
        outputfile = arg

print('Processing file: ',myfile)

opened_file=open(myfile,'r')

#The following for loop finds the length of the inputed fasta to determine the canvas width and length based on the number of sequences and the longest sequence

count = 0
longestUTR=0

for line in opened_file:
    if line[0]==">" or line[0]=="A" or line[0]=="T"or line[0]=="C"or line[0]=="G":
        count+=1
    if len(line)>longestUTR:
        longestUTR=len(line)

canvaslength=count*27.5+100
canvaswidth=longestUTR*boxwidth+550
end_of_sequence_diagrams=longestUTR*boxwidth+5

fd = EPSFile(outputfile, canvaswidth, canvaslength)


y = canvaslength


opened_file=open(myfile,'r')

#This will order the opened file by expression level. It makes a list of lists with the: mean_translation, mean_fold, sem_translation, genename, sequence. It also will determine the minimum and maximum x value for making the axis

ordered_constructs=[]

#we are starting with a min_x and a max_x that are unrealistic so we can re-set them in the code
pf_min_x=10000
pf_max_x=-10000
human_min_x=10000
human_max_x=-10000
for line in opened_file:
    if line[0]==">":
        P_position=line.find("P")
        genename=line[P_position:-1]
        if genename in plasmo_data.columns:
            #mean_fold=round(fold_data[genename].mean(),2)
            pf_mean_translation=plasmo_data[genename].mean()
            pf_mean_translation=round(pf_mean_translation,2)
            pf_sem_translation=plasmo_data[genename].sem()
            pf_sem_translation=round(pf_sem_translation,2)
            if pf_mean_translation<pf_min_x:
                pf_min_x=round(pf_mean_translation-pf_sem_translation)
            if pf_mean_translation>pf_max_x:
                pf_max_x=round(pf_mean_translation+pf_sem_translation)

            #genename=genename + " : " + str(mean_fold) + "%"
        else:
            pf_mean_translation=0
            pf_sem_translation=0
            genename=genename + " : no data"
        
        if genename in human_data.columns:
            #mean_fold=round(fold_data[genename].mean(),2)
            human_mean_translation=human_data[genename].mean()
            human_mean_translation=round(human_mean_translation,2)
            human_sem_translation=human_data[genename].sem()
            human_sem_translation=round(human_sem_translation,2)
            if human_mean_translation<human_min_x:
                human_min_x=round(human_mean_translation-human_sem_translation)
                print("min" + str(human_min_x))
            if human_mean_translation>human_max_x:
                human_max_x=round(human_mean_translation+human_sem_translation)
        else:
            human_mean_translation=0
            human_sem_translation=0
            genename=genename + " : no data"


#All  postscript starts 0,0 in the lower Left corner!
    if line[0]=="A" or line[0]=="T" or line[0]=="G" or line[0]=="C":
        sequence=line[0:-1]
        ordered_constructs.append([ pf_mean_translation, pf_sem_translation, genename, sequence, human_mean_translation, human_sem_translation])
    # ordered_constructs.append([ mean_translation, mean_fold, sem_translation, genename, sequence]). This is if you want to add the fold above P60 or another value to the title


#We need the axis to have 0 in it, so if zero is not in the range we will set the min or the max to it as needed: For plasmodium data
if pf_min_x>=0:
    pf_min_x=0
    pf_y_axis_x=end_of_sequence_diagrams+20
if pf_max_x<=0:
    pf_max_x=0
    pf_y_axis_x=end_of_sequence_diagrams+250

#pf_min_x=-2  #These two LINES CAN BE CHANGED TO SET THE X-AXIS MINIMUM TO BE CONSISTENT WITH OTHER GRAPHS
#human_min_x=-2

#pf_max_x=2 #These two LINES CAN BE CHANGED TO SET THE X-AXIS MINIMUM TO BE CONSISTENT WITH OTHER GRAPHS
#human_max_x=2

pf_tick_mark_distance=round(200/((abs(pf_min_x)+abs(pf_max_x)+1)))
pf_zero_distance=pf_tick_mark_distance*(abs(pf_min_x))

if pf_min_x!=0 or pf_max_x!=0:
    pf_y_axis_x = pf_tick_mark_distance*(abs(pf_min_x))+ end_of_sequence_diagrams + 20

#We need the axis to have 0 in it, so if zero is not in the range we will set the min or the max to it as needed: For human data
if human_min_x>=0:
    human_min_x=0
    human_y_axis_x=end_of_sequence_diagrams+20
if human_max_x<=0:
    human_max_x=0
    human_y_axis_x=end_of_sequence_diagrams+250

human_tick_mark_distance=round(200/((abs(human_min_x)+abs(human_max_x)+1)))

if human_min_x!=0 or human_max_x!=0:
    human_y_axis_x = human_tick_mark_distance*(abs(human_min_x))+ end_of_sequence_diagrams + 270


#Finds the number of tick marks we will need

human_zero_distance=human_tick_mark_distance*(abs(human_min_x))


#This will draw the axis based on the values calcuted in the previous for loop

y_axis_y=canvaslength-30

make_axis(pf_min_x, pf_max_x, human_min_x, human_max_x, count, end_of_sequence_diagrams, canvaslength, pf_y_axis_x, pf_tick_mark_distance, human_y_axis_x, y_axis_y, human_tick_mark_distance)

#This orders our new list of sequences by the first value in the sub-lists
#ordered_constructs.sort(reverse=True)

#The next part of the code goes through our new list "ordered_constructs" and draws the sequences and bars etc. with the data


for construct in ordered_constructs:
    
    genename=construct[2]
    pf_mean_translation=construct[0]
    pf_sem_translation=construct[1]
    sequence=construct[3]
    human_mean_translation=construct[4]
    human_sem_translation=construct[5]
    print(construct)
    #checks if line is a header or if it's sequence
    x = boxwidth
    boxheight = 15
    #This looks at the likes at all the name lines and grabs the construct number by looking for the position of "P" then grabbing the rest
    #so if you use numbering to order your fasta file will remove the numbering
    y-=55
    addname(x,y,genename)
    
    #Below calls the two funcitons that add data bars to the graphs
    pf_add_data_bar(pf_mean_translation, pf_sem_translation, y, pf_y_axis_x, pf_tick_mark_distance)
    human_add_data_bar(human_mean_translation, human_sem_translation, y, human_y_axis_x, human_tick_mark_distance)

        #Determine the point length of the drawen sequence
    seqlength = len(sequence)
    pointseqlength=seqlength*boxwidth
    print("pre-rbg selection")
    for mycharacter in sequence:
        #	R G B setrgbcolor  Values = 0 to 1
        if mycharacter == "A":
            fd.append("0.8 setgray")
        if mycharacter == "T":
            fd.append("0.6 setgray")
        if mycharacter == "G":
            fd.append("0.8 0 0 setrgbcolor")
        if mycharacter == "C":
            fd.append("0.5 0 0 setrgbcolor")

            #Remember - the starting point x,y is the lower left corner!
        fd.append("%f %f %f %f rectfill"%(x,y,boxwidth,boxheight))
        x = x + boxwidth

        #Find all the stops
        stops=[]
    for j in range(0,seqlength):
        if sequence[j:j+3] == "TAA" or sequence[j:j+3] == "TAG" or sequence[j:j+3] == "TGA":
            stops.append(j)
    print("now have stops")
        # Find and plot the ATGs in the sequence
    for j in range(0,seqlength):
        if sequence[j:j+3] == "ATG":
            therestop=0
            if (seqlength-j)%3==0:
                x = (j+1) * boxwidth
                shade=1
                yposition=y
                startcodon(x,yposition,shade,fd,boxwidth)
                for stop in stops:
                    if stop>j and (stop-j)%3==0:
                        z = (stop-j)*boxwidth
                        uORF(x, yposition, z, fd,boxwidth)
                        therestop=1
                        break
                if therestop==0:
                    distance_from_end=seqlength-j
                    nostop(x,yposition,fd,seqlength,distance_from_end,boxwidth)
            
            elif (seqlength-j)%3==1:
                x = (j+1) * boxwidth
                shade=0.6
                yposition=y-5
                startcodon(x,yposition,shade,fd,boxwidth)
                for stop in stops:
                    if stop>j and (stop-j)%3==0:
                        z = (stop-j)*boxwidth
                        uORF(x, yposition, z, fd,boxwidth)
                        therestop=1
                        break
                if therestop==0:
                    distance_from_end=seqlength-j
                    nostop(x,yposition,fd,seqlength,distance_from_end,boxwidth)

            
            elif (seqlength-j)%3==2:
                x = (j+1) * boxwidth
                shade=0.4
                yposition=y-10
                startcodon(x,yposition,shade, fd,boxwidth)
                for stop in stops:
                    if stop>j and (stop-j)%3==0:
                        z = (stop-j)*boxwidth
                        uORF(x, yposition, z, fd,boxwidth)
                        therestop=1
                        break
                if therestop==0:
                    distance_from_end=seqlength-j
                    nostop(x,yposition,fd,seqlength,distance_from_end,boxwidth)
            else:
                continue

# Close the EPS file.
fd.close()
print("Done\n")
```
